# Markovian Dynamics and Spectral Relaxation of Metastatic Networks

**DOI:** 10.64898/2026.08.13.743956

**Authors:** David H. Margarit

## Abstract

Structural network representations of metastatic dissemination typically focus on static topology without resolving transport dynamics, relaxation timescales, or steady-state behaviour. Here, we formulate a discrete Markovian transport model on a directed higher-order network with transition rates derived from qualitative clinical affinity classes. By constructing a non-Hermitian row-stochastic transfer operator, we characterise the relaxation dynamics through its spectral decomposition. The system exhibits a fast-mixing regime characterised by a spectral gap of *γ* ≈ 0.67, corresponding to a characteristic relaxation timescale of *τ* ≈ 1.49 discrete steps, with the influence of the primary tumour origin progressively attenuated during dissemination. Convergence towards a non-equilibrium steady state (NESS) is accompanied by a reduction in Shannon entropy, concentrating probability mass within specific topological sinks. This spectral relaxation delineates two distinct dynamical regimes: early transient dissemination (*n* < *τ*), dominated by local organ-specific transition probabilities (organotropism), and the asymptotic regime (*n* > *τ*), determined increasingly by the global transport architecture of the network. Comparison with independent clinical and autopsy observations across 21 primary tumours and 23 target organs indicates that the predicted stationary distribution is consistent with the observed hierarchy of metastatic organ involvement.

## 1 Introduction

The systemic progression of metastatic cancer can be viewed as a stochastic transport process operating far from equilibrium, shaped by the interplay between local biological affinities and the global architecture of the host network [2]. Network-based approaches have traditionally represented metastatic dissemination using dyadic graphs, describing disease propagation as a sequence of independent pairwise transitions between organs.

Although these models have provided important insights into anatomical connectivity, they are inherently limited in capturing higher-order interactions generated by the simultaneous involvement of multiple secondary sites. In complex systems, such co-occurrences often give rise to collective behaviour that cannot be explained solely through pairwise interactions. To address this structural limitation, our previous work [41] introduced a higher-order hypergraph representation in which target organs are grouped according to their clinical co-occurrence across primary tumour patterns. Using canonical configuration null models [22], that study disentangled statistically significant higher-order associations from degree-driven effects, establishing the static structural topology of metastatic progression.

However, static hypergraph representations cannot resolve the temporal dynamics of probability transport, directional asymmetry, or the relaxation of the system towards a non-equilibrium steady state (NESS) [14]. In particular, a purely topological model cannot determine the timescale over which dependence on the primary tumour origin dissipates, nor can it determine how the global architecture of the host network shapes the asymptotic distribution of metastatic risk. Resolving these dynamics requires extending the static structural description to an operator-theoretic framework grounded in non-equilibrium statistical mechanics.

In this work, we formulate a discrete Markovian transport process defined over the higher-order network structure established in [41]. The dynamics are parameterised using multi-centre clinical classifications summarised in Table 1, which categorise the propensity of 21 primary tumour tissues to establish macro-metastases across 23 target organs into discrete affinity tiers (*Common*, *Occasional*, and *Rare*). These qualitative affinity levels are mapped onto dimensionless transition rates, Λ*_ij_*, representing effective transport barriers.

**Table 1:** Clinical metastatic patterns for hypergraph construction. Primary tumour sites mapped to target organs categorised by clinical dissemination propensity (Common, Occasional, and Rare).

| Primary site | Potential target organs |  |  |
| --- | --- | --- | --- |
|  | Common | Occasional | Rare |
| Head & neck | Lymph nodes | Lung, bone, liver | Brain, skin |
| Lung | Bone, brain, Lymph nodes, liver, adrenal | Pleura | Kidney, thyroid, diaphragm, peritoneum, skin, pancreas, spleen, breast |
| Kidney | Lung | Liver, bone | Brain, adrenal, skin, thyroid, pancreas, spleen, breast |
| Pancreatic | Liver, peritoneum | Lung, bone | Stomach, colon, brain |
| Bladder | Lung, bone, liver | Lymph nodes, peritoneum, adrenal | Kidney, rectum, colon, prostate, ureter, pleura, brain |
| Testicular | Lymph nodes, lung | Liver | Bone, brain |
| Melanoma | Lymph nodes, lung, liver, brain, bone | Peritoneum | Adrenal, pancreas, spleen |
| Uterine | Lung | Lymph nodes, liver, ovary | Bone, peritoneum |
| Rectal | Liver, lung | Peritoneum, bone | Brain, adrenal, ocular |
| Prostate | Bone, Lymph nodes | Lung, liver | Peritoneum, brain, adrenal, kidney, pancreas |
| Thyroid | Lung, bone | Lymph nodes | Brain, liver, peritoneum |
| Oesophageal | Lymph nodes, liver, lung | Bone | Brain, adrenal |
| Breast | Lymph nodes, bone, lung, liver | Pleura, brain, adrenal, skin | Peritoneum, pancreas, spleen, ocular |
| Stomach | Liver, peritoneum | Lung, bone, Lymph nodes | Ovary |
| Colon | Liver | Lung, peritoneum | Bone, brain, bladder, stomach, skin, ocular, spleen |
| Ovarian | Peritoneum, diaphragm, liver | Lymph nodes, pleura | Bone, brain, colon, lung, skin |
| Liver | Lung, bone, Lymph nodes, peritoneum | Adrenal | Brain |
| Cervical | Lymph nodes, lung | Bone, liver, peritoneum | Brain |
| STS (*Soft tissue sarcoma*) | Lung | Bone, liver | Brain, adrenal |
| Ocular | Liver | Lung, bone | Skin, brain |
| BS (*Bone sarcoma*) | Lung | Liver | Brain, kidney, adrenal |

Enforcing probability conservation yields a row-stochastic transfer operator, *W̃*, governing the spatio-temporal evolution of metastatic probability mass. Because metastatic dissemination is inherently directional, *W̃* is non-Hermitian, capturing the anisotropic flow of metastatic risk across the host system.

This dynamical formulation models the host system as a discrete state space in which progression is governed by spectral transport. Through spectral decomposition of *W̃*, we demonstrate that the system operates in a fast-mixing regime where the spectral gap governs the exponential decay of initial-condition dependence. This spectral relaxation delineates a two-scale dynamical hierarchy that connects local microenvironmental preferences with global network constraints: local organotropism [20, 42] governs the transient regime (*n < τ*), whereas the spectral properties of the host network dictate the asymptotic steady-state risk (*n* ≫ *τ*). As the system approaches NESS, probability mass concentrates within a restricted set of topological sinks driven by entropy contraction.

By embedding clinically calibrated transition rates into a non-Hermitian transport framework, this study extends higher-order representations of metastatic dissemination from static topology to transport dynamics. Our results indicate that late-stage metastatic patterns emerge as non-equilibrium steady states whose asymptotic structure is governed primarily by the spectral organisation of the host network rather than by the primary tumour location. The remainder of this manuscript is organised as follows. Section 2 details the construction of the row-stochastic transfer operator *W̃* and the higher-order Markovian framework [13]. Section 3 presents the spectral analysis [15], characterising the spectral gap [43], Total Variation Distance [12], and Shannon entropy reduction [52]. Section 4 compares the emergent properties of the theoretical framework with independent evidence from clinical cohorts, post-mortem autopsy registries, and genomic studies of metastatic dissemination. Section 5 explores the biological implications of topological sinks and ergodic mixing [46]. Finally, Section 6 summarises the key conclusions and outlines directions for adaptive transport modelling.

## 2 Methods: Stochastic Formalism

### 2.1 Construction of the Metastatic Transfer Operator

To describe the systemic evolution of the disease, we map the structural information of the metastatic hypergraph onto directed Markovian dynamics. This formulation treats the organism as a discrete state space where propagation kinetics are governed by a transfer operator derived from biological affinities. The system’s fundamental structure is defined by an incidence matrix *H* ∈ {0, 1}*^M×N^*, where the *M* = 23 rows represent secondary target organs (nodes) and the *N* = 21 columns represent primary tumour patterns (hyperedges). This hypergraph framework was developed utilising clinical data curated in Ref. [41] and is detailed in Table 1.

Based on the hypergraph formalism, each hyperedge *e_j_* ∈ *E* corresponds to a specific primary cancer site, encompassing secondary target organs reported in its metastatic pattern. Explicitly, the network components are defined as follows:

- The vertex set *V* = {*v*_1_*, v*_2_*, . . ., v_M_* }, corresponding to the *M* = 23 secondary target organs: *V* = {Lymph nodes, Bone, Brain, Pleura, Diaphragm, Liver, Kidney, Adrenal, Thyroid, Stomach, Colon, Peritoneum, Rectum, Prostate, Ureter, Lung, Ovary, Bladder, Skin, Pancreas, Spleen, Ocular, Breast}.
- The hyperedge set *E* = {*e*_1_*, e*_2_*, . . ., e_N_* }, corresponding to the *N* = 21 primary cancers acting as structural drivers: *E* = {Head and Neck, Lung, Kidney, Pancreatic, Bladder, Testicular, Melanoma, Uterine, Rectal, Prostate, Thyroid, Oesophageal, Breast, Stomach, Colon, Ovarian, Liver, Cervical, STS (*Soft tissue sarcoma*), Ocular, BS (*Bone sarcoma*)}.

In the binary hypergraph formalism [10]:

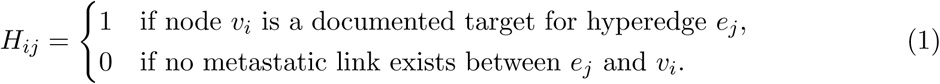

To incorporate the multi-scale and heterogeneous nature of organotropism, we define a weighted incidence matrix *H_w_* ∈ ℛ*^M×N^* via the Hadamard product [59]:

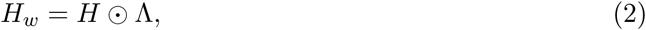

where Λ ∈ ℛ*^M×N^* is the scaling weight matrix. Its elements Λ*_ij_* represent dimensionless affinity weights encoding the relative affinity for metastatic dissemination between a primary tumour site *j* and a secondary target organ *i*.

Let *C*(*i, j*) denote the clinical classification function mapping the organ-tumour pair (*i, j*) to its qualitative affinity category. We assign the corresponding dimensionless affinity weights using a logarithmically spaced scale, with successive affinity classes separated by one order of magnitude:

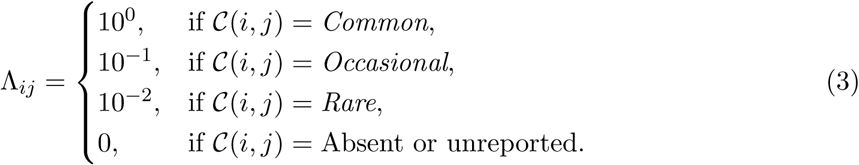

These values provide a relative weighting of the qualitative clinical categories and do not represent absolute probabilities or measured transition rates. While the qualitative clinical classifications (*Common*, *Occasional*, and *Rare*) reflect the consensus on epidemiological frequency across the established literature [1, 8, 24, 64, 65] and are systematically compiled by Margarit et al. [41], these sources do not provide explicit numerical transition rates. Consequently, any dynamical formulation requires an explicit mapping from ordinal clinical information to numerical transport weights.

Rather than introducing additional free parameters through data fitting, we adopt a parsimonious logarithmic weighting scheme by assigning successive orders of magnitude to the three affinity classes. This choice deliberately favours model parsimony over empirical overfitting, allowing the influence of higher-order network topology to be isolated from uncertainties associated with unavailable kinetic parameters. Thus, this construction preserves the ordinal hierarchy of the clinical evidence while introducing the smallest quantitative scale separation consistent with the qualitative evidence necessary to distinguish dominant, intermediate, and suppressed transport pathways.

From the perspective of statistical physics, the logarithmic spacing between successive affinity classes admits an interpretation analogous to an effective energy landscape, where larger affinity weights correspond to lower effective barriers according to the Boltzmann-type relation Δ*G_ij_* ∝ — ln Λ*_ij_* [6, 50]. We emphasise that this relation is used solely as a physical interpretation of the adopted weighting scheme rather than as an experimentally calibrated energetic model.

The assigned values therefore represent coarse-grained relative transport propensities instead of measured kinetic rate constants. Their robustness against moderate perturbations is evaluated through the Global Sensitivity Analysis presented in Sec. 3.4. Their purpose is not to reproduce absolute metastatic frequencies, but to preserve the relative ordering encoded by the underlying clinical classification.

The weighted projection between secondary organs is represented by the directed matrix *W* ∈ ℛ*^M×M^* [4]:

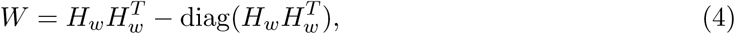

where subtraction of the diagonal removes self-loops, leaving only inter-organ interactions.

To formulate the dynamics as a discrete-time Markov process, the projection matrix *W* is row-normalised to obtain the row-stochastic metastatic transfer operator *W̃* ∈ R*^M×M^*:

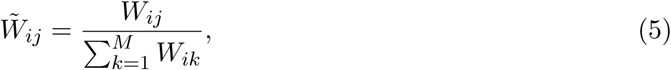

Each row of *W̃* therefore defines the conditional probability distribution over all admissible destination organs given the current occupied organ, satisfying the Markov normalisation condition [13]:

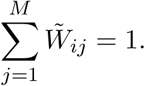

Accordingly, *W̃* defines the discrete-time Markov operator governing the evolution of metastatic probability mass over the higher-order host network. The matrices referenced by Eqs. 1, 2, 3, 4, and 5 are detailed in Appendix A of the Supplementary Material.

The resulting transport architecture is illustrated in Fig. 1. The binary incidence matrix *H* (left) defines the admissible higher-order connectivity extracted from the clinical data, whereas the corresponding transfer operator *W̃* (right) quantifies the local probability redistribution between secondary organs after row normalisation. The heterogeneous intensity pattern reflects the structural anisotropy inherited from the higher-order hypergraph, highlighting preferential dissemination pathways while excluding structurally forbidden transitions. This operator forms the basis of the spectral analysis developed in the following sections, where its eigenvalue spectrum determines relaxation dynamics, mixing behaviour, and the emergence of the non-equilibrium steady state.

**Figure 1:**
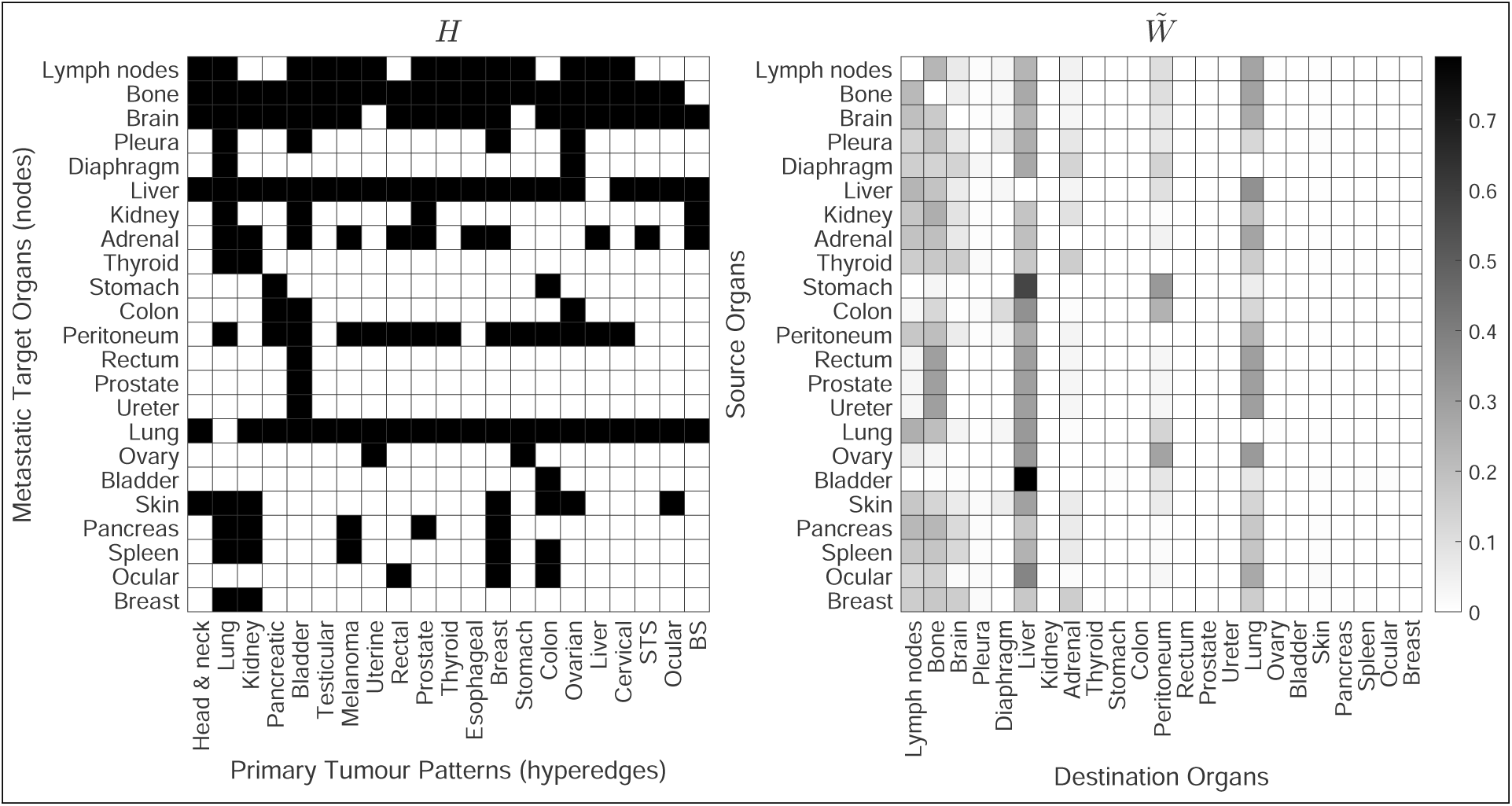
Network structural topology and local probability transport. (Left) Hypergraph incidence matrix *H* ∈ {0, 1}*^M×N^* representing the static connection landscape (*M* = 23 secondary target organs; *N* = 21 primary tumour clinical patterns). The binary mapping is shown in inverted greyscale where black cells denote an active incidence (1) and white cells indicate clinical absence (0). (Right) Projected row-stochastic metastatic transfer operator *W̃* ∈ ℛ *^M × M^* modelling local probability fluxes for discrete-time Markovian dynamics. Greyscale intensity represents the conditional transition probability from a source secondary site *i* to a destination target node *j*. The greyscale intensity is bounded within [0, 0.7] to enhance contrast, allowing low-probability pathways to remain resolvable against the structural zero background (white cells). Rows satisfy the conservation invariant ∑ *_j_ W̃_ij_* = 1.

### 2.2 Markovian Dynamics and Spectral Characterisation

The temporal evolution of the system is formalised through a discrete-time Markov chain defined over the hypergraph state space, operating under the row-stochastic transfer operator *W̃* established in Eq. 5. Each element *W̃_ij_* represents the conditional probability of a metastatic transition from source organ *i* to destination target organ *j*, satisfying the row-sum conservation invariant ∑ *_j_ W̃_ij_* = 1.

The state of the system at any discrete step *n* is described by the probability vector *p̄*(*n*) ∈ ℛ^1×*M*^, quantifying occupancy probability across secondary sites. Driven by local network flows, this distribution evolves according to the linear master equation [26], given by Eq. 6:

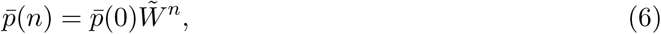

where *p̄*(0) defines the highly localised initial configuration associated with the primary tumour location [9]. Specifically, we assume that at *n* = 0, the entire probability mass is strictly concentrated at primary tumour node *i*. Mathematically, this corresponds to a Kronecker delta distribution, *p̄_j_*(0) = *δ_ji_*, establishing a deterministic, zero-entropy initial condition [63]. Eq. 6 propagates the occupancy probability mass across the network after *n* discrete transition steps. Originating from this pure state, it maps how the multi-organ colonisation likelihood shifts after *n* successive rounds of secondary dissemination governed by the stochastic transfer operator *W̃* . Given that metastatic transport pathways are directed and inherently irreversible (*W_ij_* ≠ *W_ji_*), *W̃* is a non-Hermitian operator with a generally complex eigenspectrum [5]. Beyond an algebraic property, this non-Hermiticity is the physical signature of a system operating far from thermodynamic equilibrium. Specifically, the asymmetric probability fluxes break time-reversal symmetry and violate detailed balance, establishing net directional currents across the hypergraph. Consequently, the stationary state 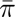 does not represent a passive equilibrium distribution, but a dynamic Non-Equilibrium Steady State (NESS) maintained by active transport kinetics. Furthermore, the global connectivity of the empirical metastatic backbone ensures that the operator is irreducible and aperiodic. Under these topological conditions, the Perron-Frobenius theorem [48] guarantees a unique, non-degenerate leading real eigenvalue *λ*_1_ = 1. Consequently, the dynamical trajectory of *p̄*(*n*) can be analytically expanded via spectral decomposition [54], naturally partitioning the global network kinetics into distinct physical regimes:

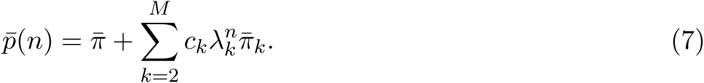

In Eq. 7, vector 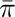 denotes the unique stationary probability of the Markov chain, defined as the leading left eigenvector satisfying:

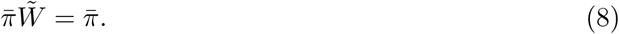

The operator *W̃* does not include absorbing states (such as organ failure or mortality).

This keeps the transition matrix ergodic, allowing the model to capture the baseline topological spread before incorporating clinical interventions or physical boundaries. Biologically, vector 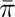 in Eq. 8 represents the steady-state distribution of metastatic flux. It functions as a global attractor dictating the intrinsic, long-term vulnerability of each secondary organ, mapping how probability mass redistributes once initial out-of-equilibrium constraints dissolve. Accordingly, spectral decomposition in Eq. 7 splits the timeline of disease into two physical phases: an early transient regime where primary tumour identity and local affinities dictate dissemination routes, and a late asymptotic regime where initial constraints dissolve, leaving host network anatomy in control.

The remaining summation in Eq. 7 constitutes the transient component of dynamics, where *k* ∈ {2*, . . ., M* } indexes non-leading relaxation modes. Here, *λ_k_* represents the *k*-th eigenvalue sorted in descending order of absolute magnitude (1 = *λ*_1_ *>* |*λ*_2_| ≥ · · · ≥ |*λ_M_* |), 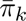 denotes its associated left eigenvector, and scalar *c_k_* ∈ R represents spectral expansion coefficients determined by projecting initial configuration *p̄*(0) onto the biorthogonal basis of the system[43].

As explored under asymptotic limits in Appendix B of the Supplementary Material, the approach to equilibrium becomes dominated by the slowest-decaying transient mode, simplifying the trajectory to 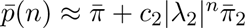. The relaxation velocity is governed by the spectral gap [43],

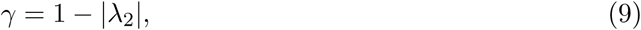

which inversely defines the characteristic relaxation timescale *τ* :

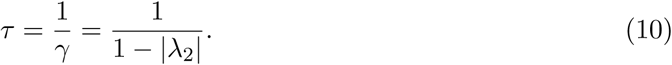

Importantly, *τ* is not an empirical fitting parameter, but an emergent physical timescale intrinsically dictated by the spectral gap of transfer operator *W̃* . While the asymptotic logarithmic mixing time is formally defined as *τ*_log_ = 1*/* ln(1*/*|*λ*_2_|) ≈ 0.90 steps, *τ* serves as a conservative linear envelope for transient decay, ensuring that *τ*_log_ *< τ*^1^.

The parameter *τ* characterises the temporal threshold of memory, establishing the characteristic scale (measured in discrete steps) required for the non-equilibrium system to dissipate localised initial seeding information. Biologically, *τ* acts as a spectral relaxation time for systemic progression, quantifying the number of dissemination steps required to erase the clinical footprint of the cancer’s origin.

To globally quantify relaxation across state space, we evaluate the Total Variation Distance (TVD) [12] between time-evolving vector *p̄*(*n*) and stationary distribution 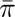:

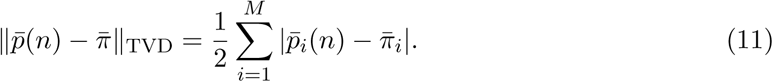

Intuitively, TVD functions as a global systemic mixing indicator, measuring remaining tissue-specific variance prior to asymptotic convergence toward the stationary distribution of the network.

### 2.3 Global Sensitivity Analysis and Structural Perturbations

To ensure that the primary network dynamical observables, namely the relaxation timescale *τ* and the stationary distribution 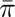, are robust spectral invariants of the host hypergraph topology rather than fine-tuned weighting artefacts, we perform a Global Sensitivity Analysis (GSA) [11] via stochastic parameter perturbation [51].

We subject the non-zero elements of the qualitative weighting matrix Λ*_ij_* to multiplicative uniform noise under a severe ±30% parameter fluctuation scale:

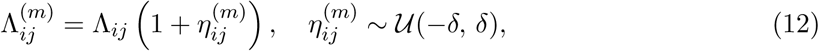

where *δ* = 0.30 bounds the relative uncertainty spanning adjacent qualitative clinical affinity tiers (*Common* = 10^0^, *Occasional* = 10*^−^*^1^, *Rare* = 10*^−^*^2^), and *m* ∈ {1*, . . ., K*} indexes independent Monte Carlo realisations (*K* = 10^4^).

For each perturbed configuration 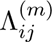 in Eq. 12, we recompute the weighted transfer operator *W̃*^(*m*)^ (Eq. 5) and extract:

- The perturbed characteristic relaxation timescale 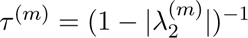.
- The perturbed stationary attractor 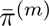 satisfying 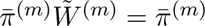.

Through Monte Carlo sampling [33] across *K* = 10^4^ realisations, we extract ensemble expectation values ℰ[*τ* ], standard deviations *σ_τ_*, and 95% empirical confidence intervals for each organ’s stationary occupancy probability.

The complete numerical pipeline and parameter perturbation protocols are formalised as pseudocode in Appendix C of the Supplementary Material. All simulations and numerical computations were performed using MATLAB version R2023b [61]. The raw implementation is available in the same reference.

## 3 Results

### 3.1 Spectral Observables and Relaxation Kinetics

The spectral analysis of the transition operator *W̃* provides a quantitative measure of the stability and efficiency of systemic metastatic spread. Within this Markovian framework, the temporal relaxation toward the stationary state is governed by the sub-dominant spectral modes [43], which dictate the rate at which the influence of the initial condition decays. Here, we formalise three key observables that bridge this operator-theoretic framework with clinical oncological progression.

First, we identify a significant spectral gap, defined in Eq. 9, as:

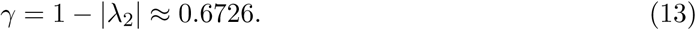

Specifically, *γ* represents the rate of information dissipation across the network. A gap of this magnitude characterises a *fast-mixing* regime, indicating that the metastatic process operates as an efficient transport mechanism where relaxation is dominated by the global architecture of the transport network. Second, this spectral gap causes the emergent relaxation timescale *τ* (Eq. 10) to collapse to a characteristic value:

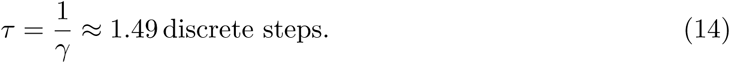

Rather than a fitted parameter, this value represents an intrinsic physical property dictated by the spectrum of the non-Hermitian operator. It implies a rapid geometric decay of the influence of the initial condition, showing that the influence of the primary tumour origin becomes strongly attenuated within fewer than two discrete transition steps.

This phenomenon is demonstrated in Fig. 2, tracking individual state-space trajectories of the 23 target organs from a primary breast seed. The rapid early flattening highlights how transient organotropism is exhausted. While the characteristic memory scale is *τ* ≈ 1.49 steps, the system requires *n* ≈ 2*τ* steps to achieve full dynamical relaxation. The vertical dashed line highlights the critical regime at *n* = 3, marking the boundary where more than 96% of initial tissue-specific variance has dissipated (|*λ*_2_|^3^ ≈ 0.035) before giving way to global ergodic convergence toward global network attractors defined by steady-state geometry 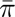 [46].

**Figure 2:**
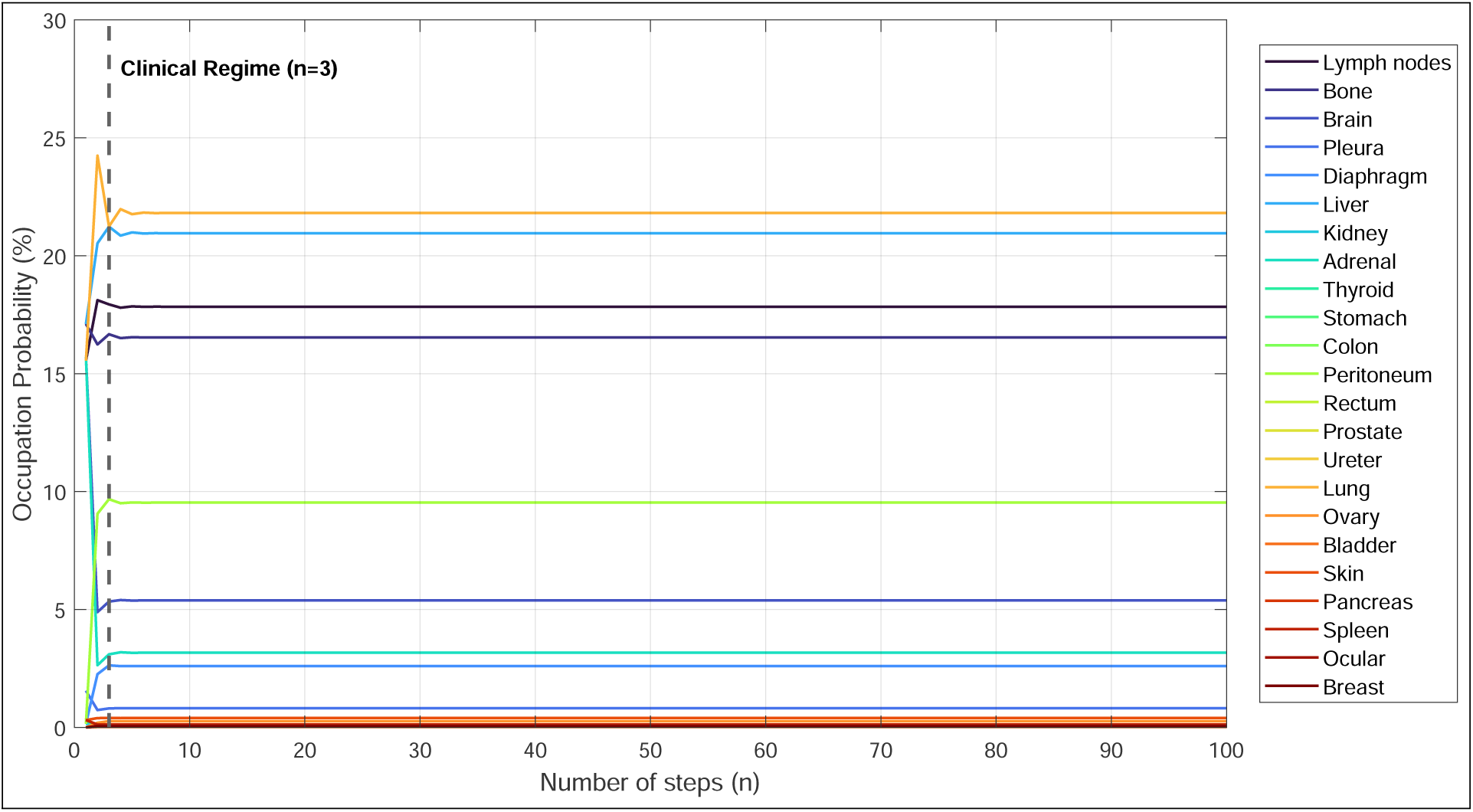
Temporal evolution of organ occupancy probabilities *p̄_i_*(*n*) initialised from a primary breast seed. Curves illustrate the trajectories of the 23 target organs towards the stationary distribution across discrete transition steps *n*. The vertical dashed line marks the reference point at *n* = 3 (≈ 2*τ*), where the transient regime gives way to asymptotic relaxation. Although the characteristic relaxation timescale is *τ* ≈ 1.49 steps, more than 96% of the initial tissue-specific variation has decayed by *n* = 3 (|*λ*_2_|^3^ ≈ 0.035). The rapid convergence is determined by the magnitude of the subdominant eigenvalue |*λ*_2_| (spectral gap *γ* ≈ 0.67), illustrating the rapid attenuation of the influence of the primary tumour origin during the earliest stages of the transport dynamics.

To globally quantify this contraction across the network, Fig. 3 plots the geometric decay of the Total Variation Distance (TVD) between the time-evolving probability vector *p̄*(*n*) and stationary distribution 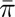 (Eq. 11).

**Figure 3:**
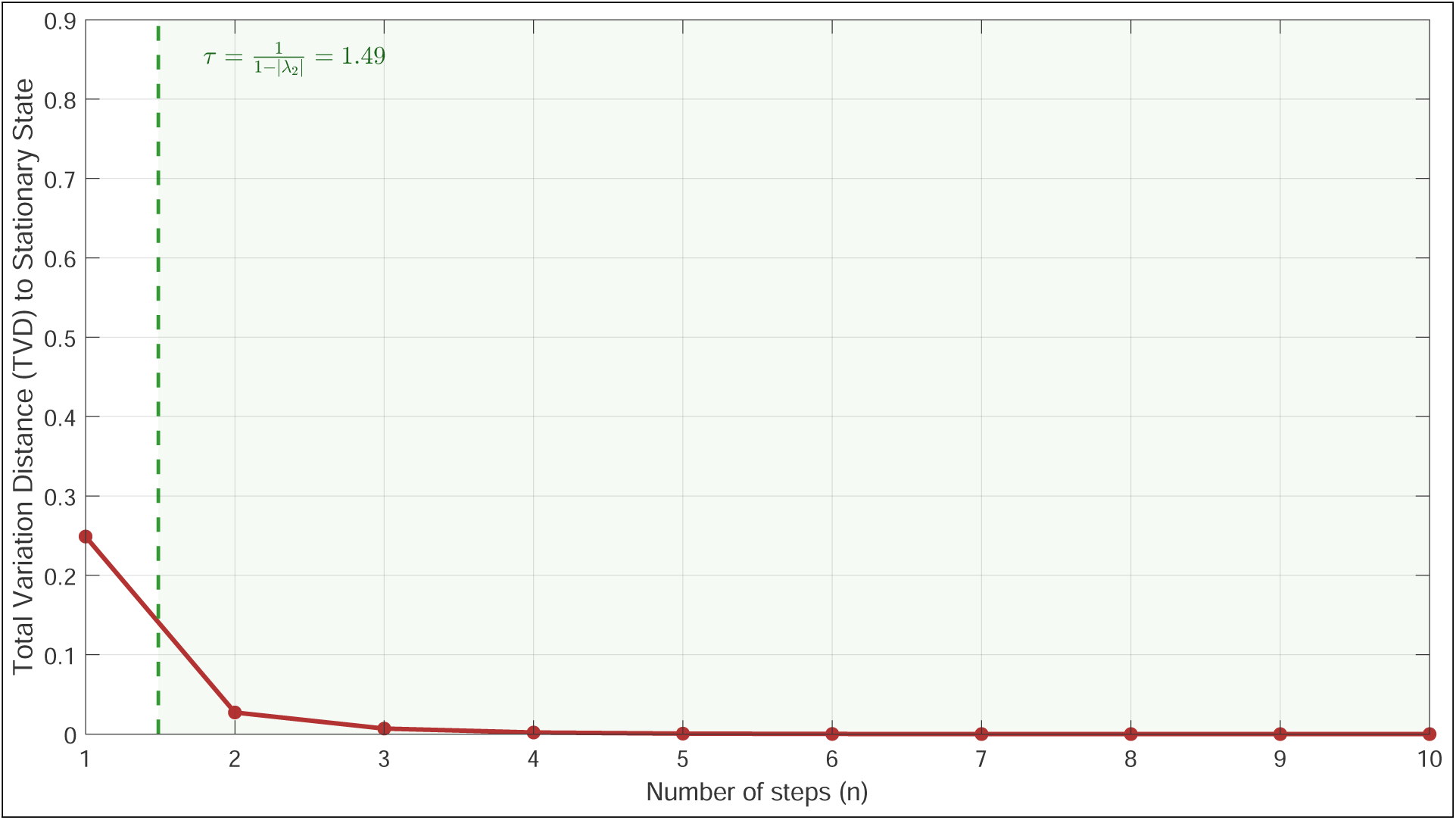
Asymptotic memory decay quantified via Total Variation Distance (TVD). The curve tracks global geometric convergence of transient occupancy vector *p̄*(*n*) toward invariant steady-state distribution 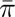 across discrete steps *n* (Eq. 11). The green dashed line marks theoretical relaxation scale *τ* = (1 − |*λ*_2_|)*^−^*^1^ ≈ 1.49 steps, serving as the formal boundary for the onset of the shaded ergodic mixing regime (*n > τ*). As TVD approaches zero, the system systematically erases the influence of the initial condition becomes negligible.

As plotted in Fig. 3, the green dashed line marks the theoretical threshold *τ* ≈ 1.49; beyond this boundary (*n > τ*), the system enters the ergodic mixing regime where the dynamics become increasingly dominated by the global architecture of the transport network rather than by the initial condition. This clean geometric decay demonstrates that the historical origin footprint is attenuated within fewer than two discrete transitions, providing a dynamical interpretation of the classical *seed-and-soil* hypothesis during advanced stages of metastatic dissemination. While initial seeding depends on specific tumour affinities (Λ), rapid spectral relaxation drives the probability distribution towards a common stationary state [20, 42].

Finally, to quantify the compartmentalisation and predictability of long-term metastatic risk, we evaluate the asymptotic Shannon entropy [52]:

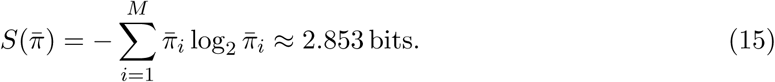

Contrasting with the maximum possible entropy for a uniform distribution over *M* = 23 secondary organs (*S*_max_ = log_2_(23) ≈ 4.524 bits), the substantial entropy reduction:

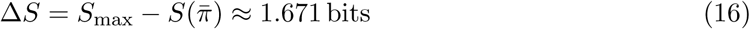

quantifies the system’s collapse toward a localised subset of organs, indicating that metastatic spread evolves as a directed transport process in which probability is channelled by the higher-order network backbone towards a restricted set of topological sinks.

### 3.2 Progression Dynamics and Ergodicity: From Local Specificity to Asymptotic Convergence

To evaluate the transition from local biology-driven to topology-driven dynamics, we tracked the evolution of the spatial probability distribution governed by the master equation (Eq. 6), initiating trajectories from distinct primary sources: lung and breast seeds (Table 2).

**Table 2:** Metastatic progression dynamics. Comparison of occupancy probabilities between specific primary seeds in transient (*n* = 3) and stationary (*n* = 100) regimes.

| Target Organ | Probability (%) |  |  |
| --- | --- | --- | --- |
| | Lung Seed ($n = 3$) | Breast Seed ($n = 3$) | Stationary State ($\bar{\pi}$) |
| Lymph nodes | 18.477 | 17.938 | 17.839 |
| Lung | 19.748 | 21.242 | 21.813 |
| Liver | 22.384 | 21.232 | 20.955 |
| Bone | 16.681 | 16.672 | 16.538 |
| Peritoneum | 9.795 | 9.678 | 9.533 |
| Brain | 5.193 | 5.337 | 5.384 |
| Ovary | 0.259 | 0.267 | 0.262 |
| Pancreas | 0.121 | 0.121 | 0.121 |
| Breast | 0.042 | 0.042 | 0.042 |
| Stomach | 0.041 | 0.040 | 0.040 |
| Prostate | 0.027 | 0.027 | 0.027 |

A detailed quantitative analysis of the dynamics reported in Table 2 highlights how out-of-equilibrium transport kinetics, facilitated by the row-stochastic operator *W̃*, progressively attenuate the influence of primary organotropism. In the early transient regime (*n* ≤ 3), the system retains subtle organ-specific clinical affinities Λ*_ij_* [41]. For instance, a primary lung seed at *n* = 3 exhibits a transient overshooting toward the liver (22.384%), temporarily exceeding local lung self-retention (19.748%). Conversely, a primary breast seed at *n* = 3 displays a symmetric co-dominance between lung (21.242%) and liver (21.232%).

Despite these initial variations, two striking dynamic properties emerge from Table 2 by *n* = 3:

- High-capacity hub convergence: The top four anatomical sinks (lymph nodes, lung, liver, and bone) collectively absorb over 76.8% of total probability mass regardless of origin, maintaining occupancy values within 1.5% of their ultimate asymptotic limits (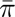).
- Instantaneous topological suppression of minor sinks: Secondary and tertiary organs with low spectral capacity (e.g., ovary, pancreas, breast, stomach, and prostate) exhibit near-instantaneous convergence. For instance, pancreatic occupancy reaches exactly 0.121% at *n* = 3 for both seeds, matching its stationary state 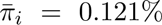 up to three decimal places. Similarly, prostate (0.027%) and breast (0.042%) occupancies show complete invariance to initial boundary conditions within *n* = 3 steps.

This rapid erosion of initial organotropism is visually summarised in Fig. 4, which compares the transient occupation vectors *p̄*(3) for lung and breast seeds against the asymptotic distribution *p̄*(100) (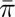).

**Figure 4:**
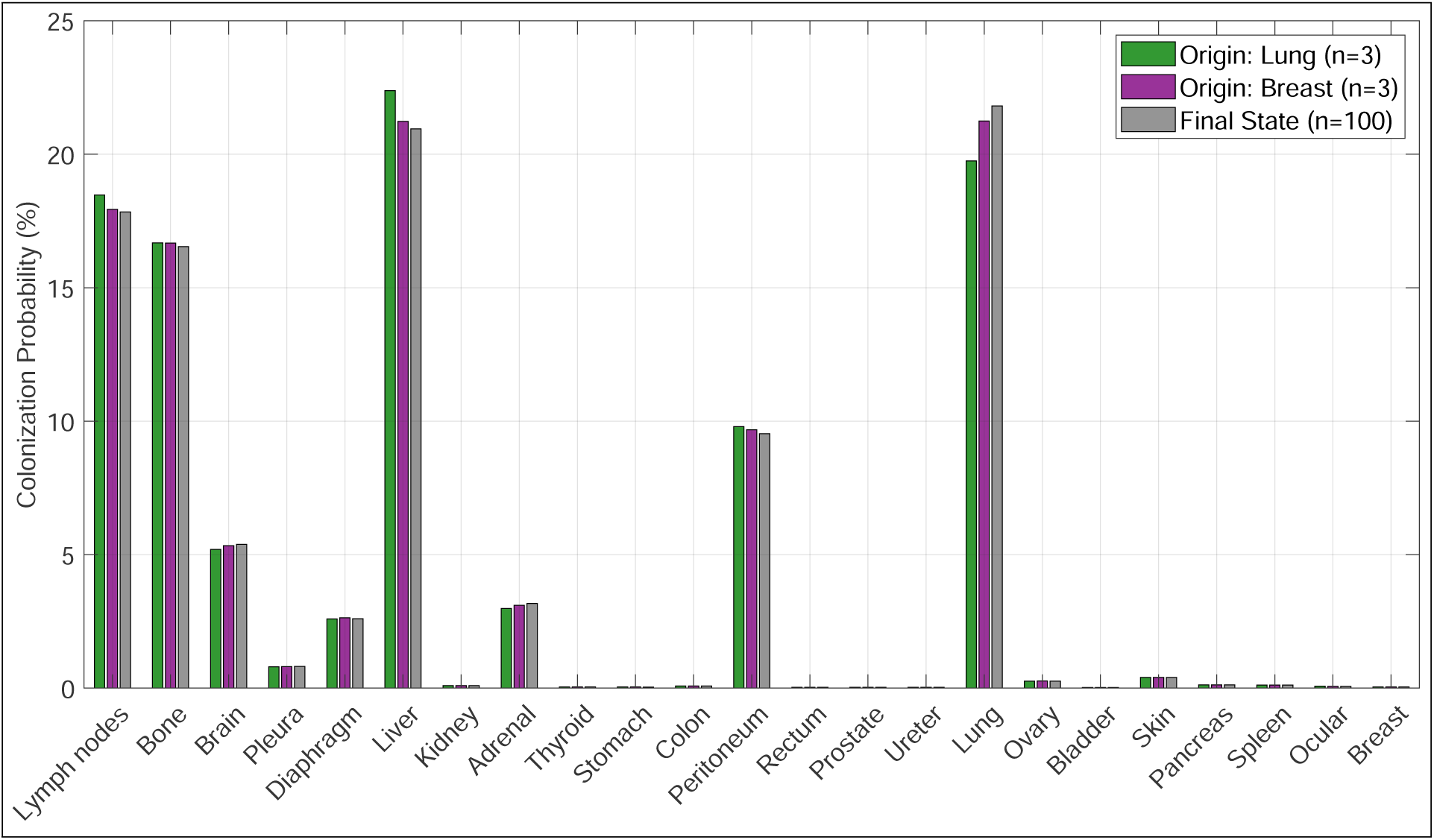
Organ-specific occupation probabilities during spectral relaxation. The bar plot compares colonisation probability profiles across 23 target organs for primary tumours originating in lung (green) and breast (violet) at *n* = 3, alongside the asymptotic non-equilibrium steady state at *n* = 100 (grey). By *n* = 3, initial differences associated with organ-specific transition probabilities have largely disappeared, illustrating rapid ergodic convergence [46] toward topological sinks.

While our previous work isolated a static structural backbone by filtering nodal degree biases [41], this dynamic formulation demonstrates that the backbone acts as a high-conductance transport pathway for probabilistic flux. Because the filtered network is strongly connected, the row-stochastic operator *W̃* is irreducible and aperiodic. By the Perron-Frobenius theorem [48], *W̃* possesses a unique leading eigenvalue *λ*_1_ = 1, with all other eigenvalues bounded strictly within the unit circle (|*λ_i_*| *<* 1).

This spectral constraint guarantees that as *n* → ∞, the system relaxes toward a unique stationary regime dictated by the leading left eigenvector 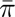. Any localised perturbations introduced by the initial configuration *p̄*(0) are exponentially suppressed at a rate governed by the spectral gap *γ*, transforming the probability distribution from one dominated by tissue-specific affinities to one governed by the stationary structure of the transport network.

### 3.3 The Hierarchy of Systemic Vulnerability

The stationary distribution 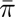 allows objective ranking of organs based on colonisation vulnerability (Table 3), revealing dominant topological sinks acting as global attractors across the metastatic hypergraph backbone. This systemic susceptibility hierarchy is graphically summarised in Fig. 5.

**Figure 5:**
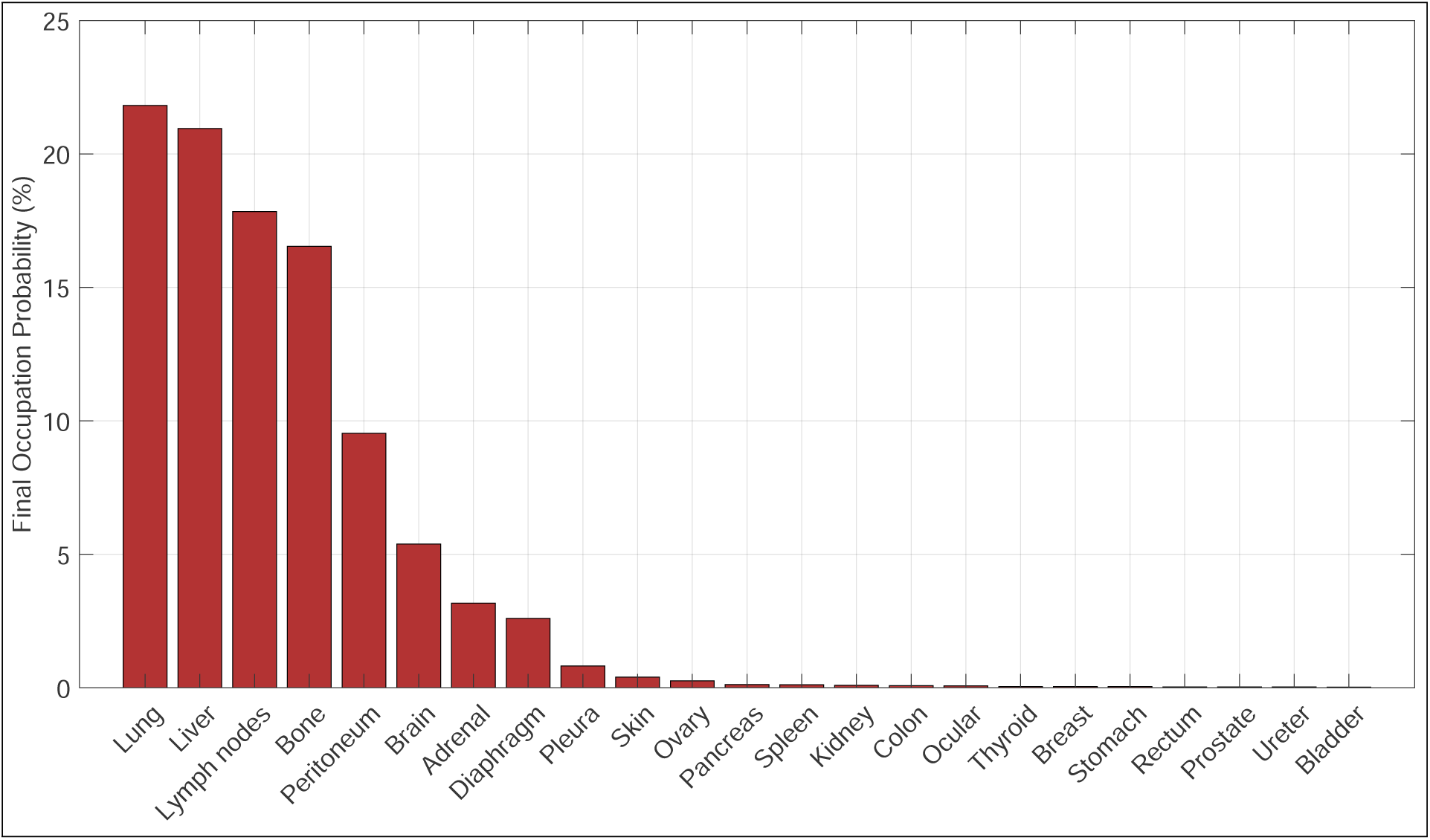
Stationary probability distribution 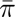 and systemic vulnerability hierarchy. The bar chart ranks 23 secondary target organs according to asymptotic occupancy probabilities from leading eigenvector of *W̃* (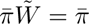 as *n* → ∞). Prominent peaks at lung (21.813%) and liver (20.955%) identify them as the dominant topological sinks of the transport network.

**Table 3:** Systemic vulnerability hierarchy. Stationary occupancy probabilities (%) representing the topological signature of metastatic risk.

| Organ | Prob. (%) | Organ | Prob. (%) |
| --- | --- | --- | --- |
| Lung | 21.813 | Pancreas | 0.121 |
| Liver | 20.955 | Spleen | 0.115 |
| Lymph nodes | 17.839 | Kidney | 0.091 |
| Bone | 16.538 | Colon | 0.076 |
| Peritoneum | 9.533 | Ocular | 0.067 |
| Brain | 5.384 | Thyroid | 0.042 |
| Adrenal | 3.168 | Breast | 0.042 |
| Diaphragm | 2.600 | Stomach | 0.040 |
| Pleura | 0.815 | Rectum | 0.027 |
| Skin | 0.397 | Prostate | 0.027 |
| Ovary | 0.262 | Ureter | 0.027 |
|  |  | Bladder | 0.020 |

The lung (21.813%) and liver (20.955%) emerge as primary sinks, accounting for over 42% of total stationary probability. Unlike models attributing this dominance solely to blood flow or local microenvironments, our spectral approach shows that their prominence emerges from the higher-order organisation of the transport network, providing a dynamical interpretation of the structural hubs identified in [41]. Conversely, organs like the bladder (0.020%) represent high-resistance nodes that remain weakly connected within the long-term transport dynamics.

### 3.4 Global Sensitivity Analysis and Structural Robustness

To rigorously confirm that the fast-mixing dynamics, spectral relaxation scale, and systemic vulnerability hierarchy are intrinsic structural features rather than artefacts of specific parameter choices, we performed a Global Sensitivity Analysis (GSA) using Monte Carlo simulations (*K* = 10^4^ realisations). In each iteration, transition rates Λ*_ij_* were independently perturbed with ±30% uniform noise around their baseline empirical values. As illustrated in Fig. 6, the characteristic relaxation timescale remains virtually invariant under parametric noise, yielding a mean GSA value of *τ*_GSA_ = 1.486±0.008 steps (with a 95% of confidence interval of [1.470, 1.502]) compared to the unperturbed baseline of *τ* = 1.487 steps. Throughout descriptive summaries and figure schematics, this timescale is expressed as *τ* ≈ 1.49 steps for conciseness; nevertheless, full three-decimal precision confirms that ergodic collapse consistently occurs within fewer than two discrete transition steps.

**Figure 6:**
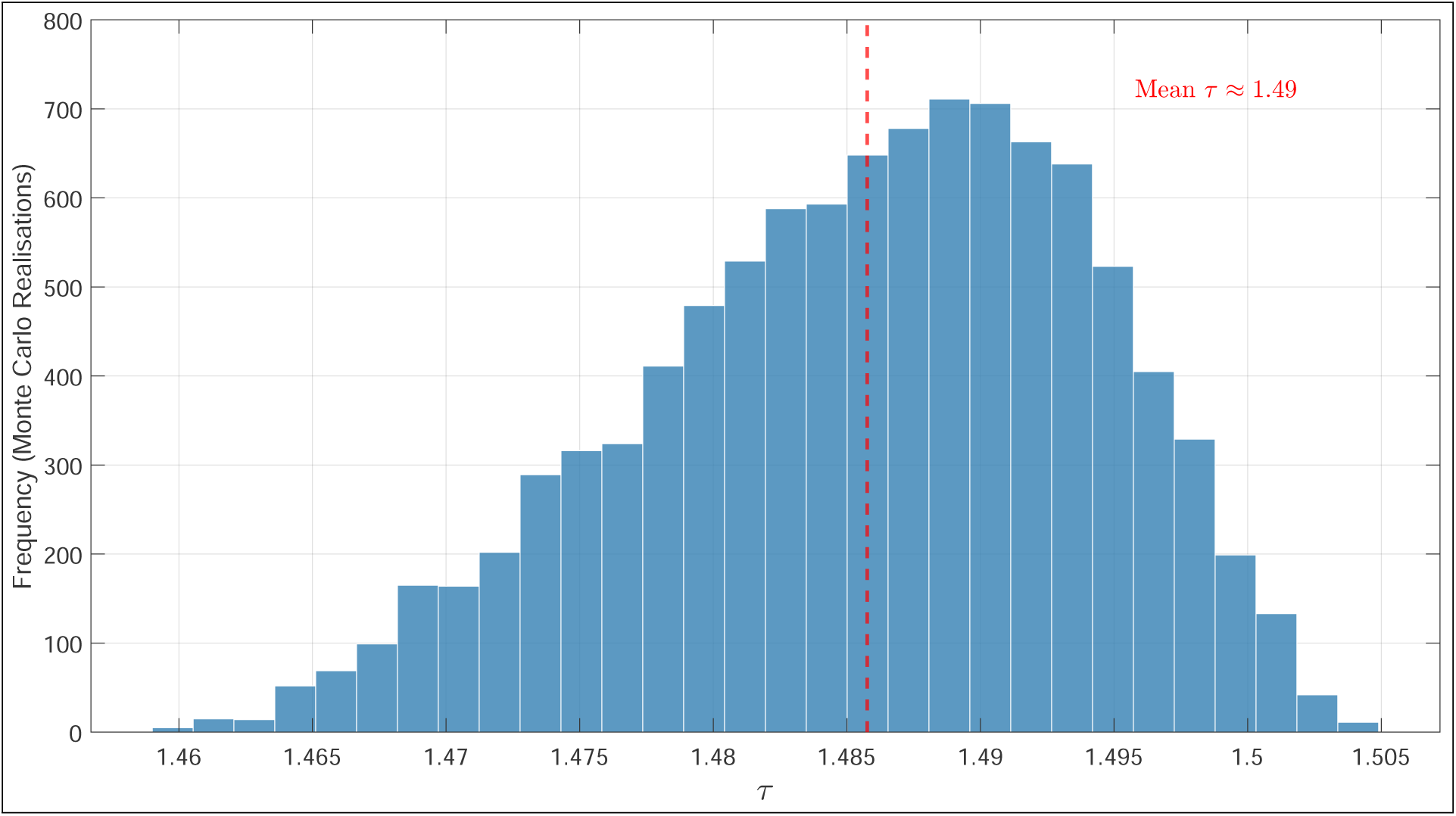
Structural robustness of the fast-mixing regime under parametric noise. istogram displays the distribution of characteristic relaxation time *τ* across *K* = 10^4^ Monte Carlo realisations subject to ±30% uniform noise perturbations applied to transition rates Λ*_ij_*. The vertical dashed line indicates the mean relaxation time (*τ*_GSA_ = 1.486 ± 0.008 steps, corresponding to *τ* ≈ 1.49 in descriptive text), demonstrating exceptional stability against parametric uncertainty.

The structural stability extends directly to individual organ occupancies, as consolidated in Table 4 and visualised via box plots in Fig. 7. The primary topological sinks exhibit remarkably tight confidence bounds (e.g., lung mean 21.786%, *σ* = 0.224%; liver mean 20.937%, *σ* = 0.137%), proving that long-term systemic vulnerability is protected by global host connectivity rather than fine-tuned local rate choices. Secondary and minor sinks display subtle positive skewness, accounted for in Table 4 via empirical non-parametric percentiles (2.5^th^ and 97.5^th^), which naturally emerges from non-linear rate propagation across multi-organ hypergraph pathways. Overall, the GSA demonstrates that while microscopic rate fluctuations slightly modulate local kinetics, the large spectral gap (*γ* ≈ 0.67) acts as a robust topological shield that preserves the asymptotic non-equilibrium steady state.

**Figure 7:**
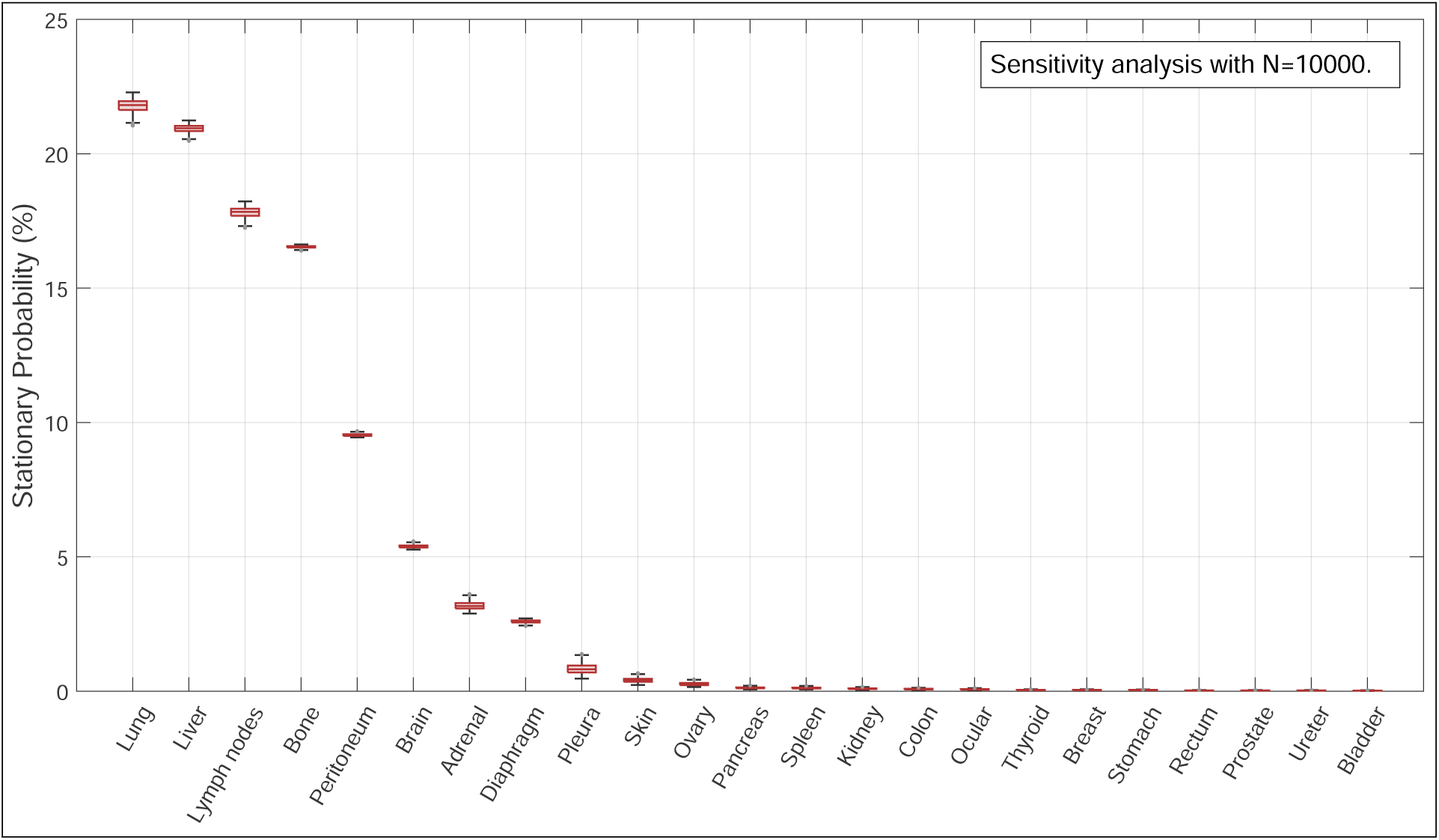
Invariance of the metastatic hierarchy under parametric uncertainty. Box plots display stationary occupancy probabilities *π̄_i_* across 23 target organs over *K* = 10^4^ realisations under ±30% uniform noise in Λ*_ij_*. Tight interquartile ranges demonstrate that universal topological sink rankings are structurally absolute.

**Table 4:** Global Sensitivity Analysis (GSA) summary. Baseline occupancy probabilities versus GSA statistics under ±30% parametric noise (Λ*_ij_* perturbed across *K* = 10^4^ realisations). *Note: The* 95% *confidence intervals correspond to empirical non-parametric percentiles (*2.5*^th^ and* 97.5*^th^), reflecting natural distribution skewness arising from non-linear rate propagation across the hypergraph. The baseline relaxation time τ* = 1.487 *steps (τ_GSA_* = 1.486 ± 0.008*) is reported with full statistical precision, corresponding to τ* ≈ 1.49 *steps in descriptive text*.

| Target Organ | Base Model (%) | GSA Mean (%) | Std. Dev. ($\sigma$) | 95% Conf. Interval |
| --- | --- | --- | --- | --- |
| Lung | 21.813 | 21.786 | 0.224 | [21.306, 22.157] |
| Liver | 20.955 | 20.937 | 0.137 | [20.643, 21.163] |
| Lymph nodes | 17.839 | 17.818 | 0.178 | [17.442, 18.117] |
| Bone | 16.538 | 16.536 | 0.041 | [16.451, 16.607] |
| Peritoneum | 9.533 | 9.535 | 0.043 | [9.462, 9.623] |
| Brain | 5.384 | 5.387 | 0.055 | [5.292, 5.501] |
| Adrenal | 3.168 | 3.183 | 0.137 | [2.955, 3.482] |
| Diaphragm | 2.600 | 2.594 | 0.055 | [2.475, 2.685] |
| Pleura | 0.815 | 0.834 | 0.183 | [0.529, 1.235] |
| Skin | 0.398 | 0.407 | 0.080 | [0.275, 0.578] |
| Ovary | 0.262 | 0.268 | 0.056 | [0.175, 0.389] |
| Pancreas | 0.121 | 0.125 | 0.029 | [0.077, 0.189] |
| Spleen | 0.116 | 0.118 | 0.028 | [0.073, 0.180] |
| Kidney | 0.091 | 0.093 | 0.022 | [0.058, 0.142] |
| Colon | 0.076 | 0.078 | 0.018 | [0.048, 0.117] |
| Ocular | 0.068 | 0.069 | 0.016 | [0.043, 0.105] |
| Thyroid | 0.043 | 0.044 | 0.010 | [0.027, 0.066] |
| Breast | 0.043 | 0.044 | 0.010 | [0.027, 0.066] |
| Stomach | 0.040 | 0.041 | 0.010 | [0.025, 0.062] |
| Rectum | 0.027 | 0.028 | 0.007 | [0.017, 0.042] |
| Prostate | 0.027 | 0.028 | 0.007 | [0.017, 0.042] |
| Ureter | 0.027 | 0.028 | 0.007 | [0.017, 0.042] |
| Bladder | 0.020 | 0.021 | 0.005 | [0.013, 0.031] |
| **Relaxation Time ($\tau$)** | **1.487 steps** | **1.486 steps** | **0.008** | **[1.470, 1.502]** |

## 4 Clinical Consistency and Empirical Evidence

The biological relevance of the proposed dynamical framework was examined by comparing its emergent properties with independent clinical observations from population-based registries, autopsy studies, and genomic analyses of metastatic progression. We distinguish between quantitative agreement with observed metastatic distributions and biological consistency with established mechanisms of dissemination. Since the model is not fitted to organ-specific prevalence data, these comparisons provide an assessment of the extent to which host-network architecture captures clinically relevant features of systemic metastatic organisation.

### 4.1 Ergodic Convergence and Terminal Sinks: Agreement with Clinical Registries and Autopsy Data

Large autopsy series and population-based cancer registries, including the NCI Surveillance, Epidemiology, and End Results (SEER) Program [23], consistently show that, despite substantial differences in primary tumour histology, advanced metastatic disease is dominated by a limited set of anatomical compartments, particularly the lung, liver, bone, and lymphatic system [31, 34, 56, 57, 65, 67]. Together, these organs account for most metastatic burden across diverse cancer types.

Remarkably, the stationary distribution predicted by our transfer operator reproduces this same hierarchy, assigning 21.81% of the probability mass to the lung, 20.96% to the liver, 17.84% to lymph nodes, and 16.54% to bone, jointly accounting for 77.15% of the asymptotic occupation probability.

Because the transfer operator is constructed solely from higher-order connectivity and qualitative affinity classes, without incorporating the observed prevalence of metastases at individual organs, this agreement represents an independent consistency check of the model rather than a directly fitted outcome. The ability of the model to recover the clinically observed metastatic hierarchy suggests that higher-order host-network topology encodes relevant structural constraints underlying systemic metastatic dissemination.

### 4.2 Spectral Memory Dissipation and Clinical Patterns of Oligo- and Polymetastatic Disease

The characteristic relaxation timescale (*τ* ≈ 1.49 steps), derived from the dominant spectral gap *γ* = 1 − |*λ*_2_| ≈ 0.6726 (Eq. 13), provides a quantitative framework for interpreting the transition between localised oligometastatic dissemination and widespread polymetastatic involvement observed in clinical oncology [27, 45].

In the early transient regime (*n* = 1), the state occupancy distribution *p̄*(1) remains heavily weighted by primary tumour organotropisms (Λ*_ij_*). This regime is compatible with the clinically observed window of organ-confined or oligometastatic spread, where local ablative interventions (such as stereotactic ablative radiotherapy evaluated in the landmark SABR-COMET trials) achieve long-term disease control by targeting histology-dependent, predictable seeding sites [45]. However, for dissemination steps beyond the relaxation timescale (*n* ≥ 2, with *n > τ*), the corresponding contribution of the leading non-stationary mode is reduced by approximately 89% (with residual memory |*λ*_2_|^2^ ≈ 0.107).

This rapid spectral mixing provides a possible dynamical interpretation of why advanced metastatic disease is frequently associated with broader, less site-specific dissemination patterns. As dissemination steps accumulate, primary-site preferences become progressively attenuated, resulting in an increased contribution of global host-network structure to metastatic occupancy profiles [28, 37].

### 4.3 Secondary Cascades and Biological Consistency with Multi-Step Dissemination

A notable feature of the multi-step Markovian model is the progressive increase in stationary occupancy of downstream secondary sinks, most notably the peritoneum 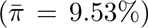, brain 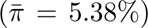, and adrenal glands 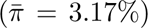 [8, 24, 64]. For the majority of primary solid tumours, these sites exhibit comparatively limited direct seeding during the initial dissemination wave (*n* = 1) [19]. Their increasing contribution at later steps therefore reflects a network-level signature of sequential dissemination, in which organs that are not dominant initial targets can acquire greater occupancy through successive transitions across the metastatic network.

This behaviour is biologically consistent with experimental and genomic evidence indicating that metastatic progression can involve secondary colonisation and cross-seeding between already established metastatic sites [55]. Genomic phylogenetic analyses in human cohorts further support the occurrence of sequential dissemination events, showing that visceral sites, particularly the liver and lung as major vascular receiving compartments, can subsequently participate in further metastatic spread [30, 47]. In this context, the present model provides a network-level representation of a process in which previously colonised organs may contribute indirectly to subsequent dissemination, consistent with observations of metastatic diversification reported in human studies [18, 35]. Importantly, this correspondence should be interpreted as biological consistency rather than direct mechanistic validation of clonal cross-seeding, since the present model does not explicitly represent tumour clones, phylogenetic relationships, or organ-to-organ cellular transfer.

## 5 Discussion: The Physics of Systemic Convergence

The results show that the higher-order organisation of the host network plays a central role in shaping the long-term dynamics of metastatic dissemination. By extending a static higher-order network representation into an operator-theoretic transport framework, we characterise how the spectral properties of the transfer operator *W̃* govern relaxation dynamics and the emergence of a non-equilibrium steady state. In the following sections, we discuss the physical interpretation of the characteristic relaxation timescale (*τ* ≈ 1.49) and its implications for metastatic organotropism, systemic transport, and host-driven constraints on disease progression.

### 5.1 Beyond the Seed-and-Soil: Spectral Relaxation and Ergodicity

Our results provide a dynamical perspective on the classical *seed-and-soil* hypothesis [20, 42]. Organ-specific dissemination has traditionally been attributed to biochemical affinities between circulating tumour cells and permissive microenvironments [49]. While our previous higher-order hypergraph model represented these affinities through statistically validated structural associations [41], the present framework incorporates them into a dynamical transport process through the transition rates Λ*_ij_*. The resulting spectral structure indicates that the large spectral gap (*γ* ≈ 0.67) promotes rapid convergence toward the stationary regime, progressively reducing the dependence of the global occupancy distribution on the initial primary site.

The progression dynamics reported in Table 2 illustrate this relaxation. After three transition steps (*n* = 3), distributions originating from different primary tumours still retain measurable differences, but already approach the stationary profile. For a primary lung tumour, the transient distribution exhibits a higher probability for liver involvement (22.384%) than for lung retention (19.748%), whereas a primary breast tumour produces an almost symmetric distribution between lung (21.242%) and liver (21.232%). Because *n* = 3 exceeds twice the characteristic relaxation time (*τ* ≈ 1.49), the dominant occupancies are already close to their stationary values, with deviations of approximately 1.5%. By *n* = 100, both trajectories converge to the same asymptotic ranking, dominated by lung (21.813%), liver (20.955%), lymph nodes (17.839%), and bone (16.538%). Thus, the model distinguishes between an early regime in which primary-site affinities strongly influence dissemination and a later regime in which global host-network structure increasingly determines organ occupancy.

The discrete variable *n* should therefore be interpreted as a topological time index rather than as chronological time. Each transition represents a coarse-grained dissemination event that may encompass processes such as intravasation, vascular transport, extravasation, and establishment within a secondary organ [25]. This interpretation should not be taken as a direct mapping onto clinical staging. In particular, *n* = 0, *n* = 1, and *n* ≥ 2 do not correspond formally to M0, M1a, and later TNM (*Tumour-Node-Metastasis*) categories, respectively. Rather, the increasing value of *n* provides a network-level representation of successive dissemination events that can be qualitatively compared with the progression from limited to multi-organ metastatic disease described in clinical classifications [3, 44]. Within this framework, the characteristic relaxation time (*τ* ≈ 1.49) indicates that the influence of the initial primary site becomes substantially attenuated after fewer than two effective dissemination transitions.

The transition rates Λ*_ij_* also admit a useful effective-energy interpretation. Under a Boltzmann-type coarse-grained description [6], transition probabilities may be related to activation barriers through Λ*_ij_* ∝ *e^−β^*^Δ*G*^*^ij^*, or equivalently, Δ*G_ij_* ∝ − ln Λ*_ij_*, where *β* = (*k_B_T*)*^−^*^1^ is absorbed into the proportionality constant [50]. Within the present framework, however, this correspondence is interpretative rather than thermodynamic: Δ*G_ij_* is not independently estimated and the affinity classes are not derived from measured free energies. The analogy instead provides a physical interpretation of the hierarchy of transition rates as representing progressively less favourable effective transport pathways. Consistent with the Total Variation Distance decay shown in Fig. 3, the resulting dynamics become increasingly governed by the spectral properties of the network rather than by the initial condition. This provides a possible mechanistic interpretation for the emergence of similar long-term organ occupancy patterns across biologically heterogeneous primary tumours.

The rapid convergence observed for *n > τ* characterises the behaviour of the idealised transport network in the absence of absorbing states, therapeutic interventions, or explicit time-dependent remodelling of the host architecture. These processes are biologically relevant and may interrupt or modify metastatic trajectories [17, 58]. Their exclusion here is therefore a modelling assumption rather than a claim that real metastatic progression is unbounded or unaffected by clinical intervention. Under this controlled setting, the model isolates the contribution of network connectivity to the redistribution of metastatic probability and provides a reference against which more realistic adaptive or intervention-dependent dynamics can be developed.

### 5.2 Topological Sinks and the Channelling of Metastatic Flux

The stationary distribution identifies the lung (21.813%) and liver (20.955%) as the dominant topological sinks of the higher-order transport network, together accounting for 42.77% of the stationary probability mass (Table 3). From the perspective of non-equilibrium statistical mechanics, these organs emerge as preferential accumulation sites from the combined effect of higher-order connectivity and the encoded transition affinities. Thus, their prominence in the stationary regime is not determined by a single local interaction, but by how local transition preferences are embedded within the global organisation of the transport network. The stationary probability vector 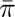 shown in Fig. 5 therefore provides a quantitative measure of the long-term occupancy propensity of each organ within the modelled transport process.

The reduction in Shannon entropy, *S*(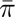) ≈ 2.853 bits (Eq. 15), relative to *S*_max_ = log_2_(23) ≈ 4.524 bits (Eq. 16), provides an information-theoretic measure of this concentration [52]. The corresponding entropy deficit, Δ*S* = *S*_max_ − *S* ≈ 1.671 bits, indicates that the stationary dynamics explore a substantially smaller effective state space than would be expected for a uniform random walk. Equivalently, the entropy difference corresponds to an effective reduction in accessible states by a factor of 2^Δ*S*^ = 2^1.671^ ≈ 3.18. Rather than distributing probability uniformly across all *M* = 23 organs, the higher-order transport architecture channels probability toward a restricted subset of network states, producing a concentrated non-equilibrium stationary distribution.

These preferential transport patterns emerge from the statistically validated higher-order backbone identified in our previous work using configuration-based null models [41]. By controlling for degree-driven effects in the observed co-occurrence data, that framework isolated higher-order associations that could not be explained by marginal connectivity alone. The present dynamical analysis extends this result by showing that these structural relationships generate persistent differences in long-term transport occupancy, with probability progressively concentrated in the principal sinks of the network. In this sense, the present framework connects the previously identified source–sink organisation (*V_S_* → *V_T_*) with an explicit dynamical description of metastatic flux.

Lymph nodes (17.839%) provide a complementary example of this network organisation. Beyond their role as early dissemination sites, they retain a substantial fraction of the stationary probability and therefore act as persistent redistribution compartments within the modelled transport process. Because the characteristic relaxation time (*τ* ≈ 1.49) indicates progressive attenuation of primary-site dependence after a small number of dissemination transitions, the relative importance of these host-level transport structures increases as the system approaches its stationary regime. From a clinical perspective, this result suggests that host-network organisation may provide complementary information to primary tumour biology when interpreting advanced polymetastatic dissemination.

## 6 Conclusions

In this work, we extended a static higher-order topological description into a dynamical Markovian framework to investigate metastatic dissemination as a directed transport process shaped by multi-organ network architecture. By separating biological affinities from structural constraints through the directed transition operator *W̃*, and assessing robustness through Global Sensitivity Analysis (*K* = 10^4^ Monte Carlo realisations under ±30% parametric perturbations), we established a quantitative framework for describing the relaxation, asymptotic organisation, and structural robustness of metastatic transport.

### 6.1 Principal Findings

Table 5 summarises the principal quantitative findings of this study together with their physical and biological interpretation. Collectively, these results describe a progression from rapid spectral relaxation and convergence to a concentrated stationary distribution, while also demonstrating robustness to parameter perturbations and consistency with independent clinical observations. The table is intended to provide a concise overview of the main results before discussing their broader implications and limitations.

**Table 5:**
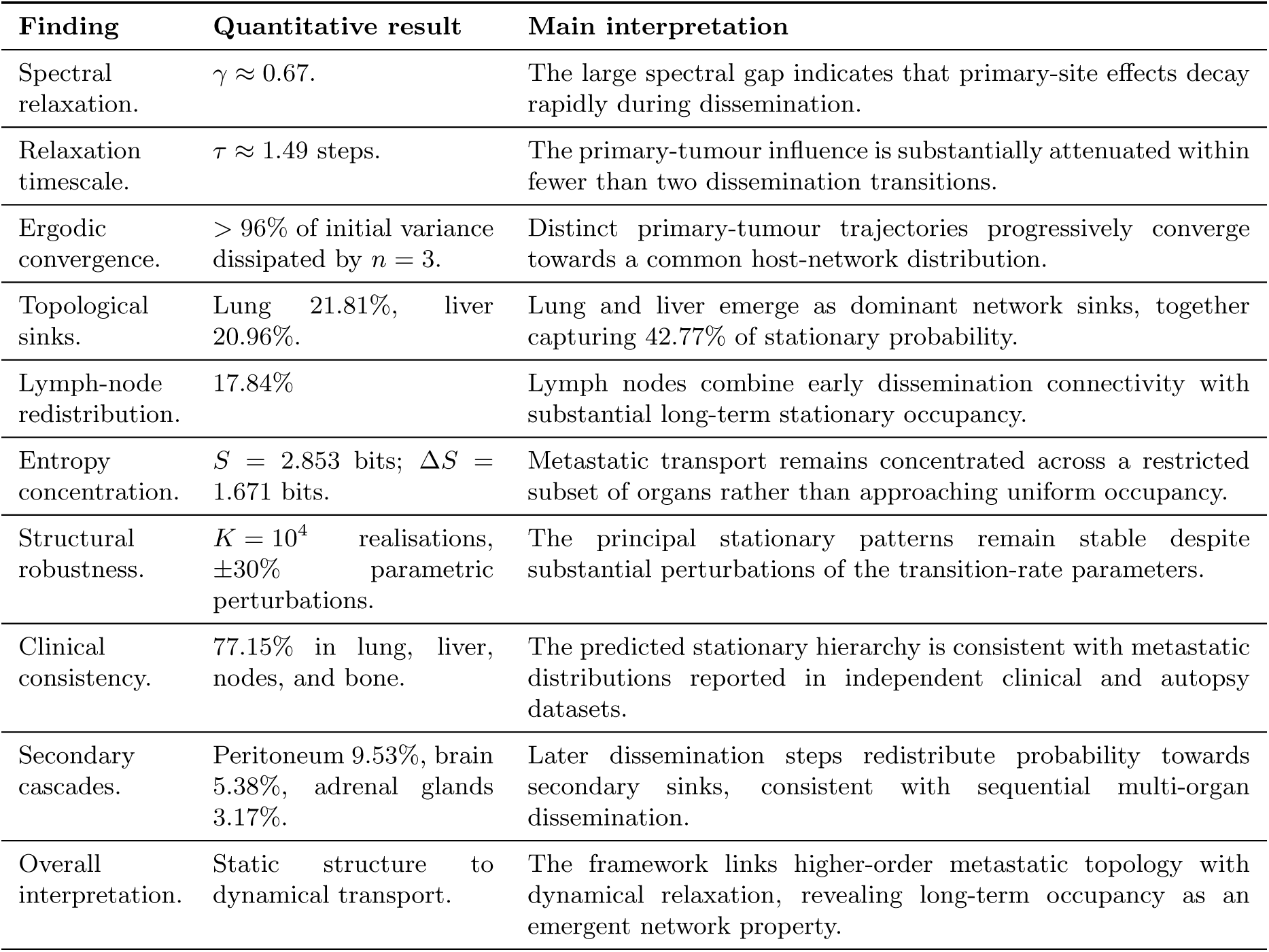
Summary of the principal findings and their interpretation.

Collectively, these findings provide the quantitative basis for the broader conceptual interpretation developed below, beginning with the transition from the static higher-order framework to the present dynamical formulation.

### 6.2 Comparative Analysis: From Static Structure to Dynamical Ergodicity

This dynamical approach marks a conceptual shift from our previous structural framework [41]. In our earlier work, we utilised a canonical hypergraph configuration model (via Metropolis-Hastings MCMC sampling) to filter out degree-driven popularity biases and isolate a statistically validated structural backbone, establishing a functional partition between source (*V_S_*) and sink (*V_T_*) organ cohorts [41]. However, that formulation remained strictly static, mapping permissible topological boundaries without revealing temporal kinetics, transport efficiency, directional relaxation, or asymptotic limits.

The present study resolves this limitation by extending that static backbone into a dynamical transport framework under out-of-equilibrium physical conditions. Mapping qualitative clinical affinities onto dimensionless transition rates Λ*_ij_* transforms passive co-occurrence associations into a directed probability flux. While the static model identified *where* disease could spread, this spectral framework establishes *how* and *how fast* it spreads. It uncovers a fast-mixing regime governed by a significant spectral gap (*γ* ≈ 0.67, *τ* ≈ 1.49), where host architecture drives trajectories originating from different primary tumours towards a common stationary distribution into a universal ergodic steady state 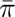 [68]. Consequently, we shift the analytical focus from static multi-organ pattern mapping to the quantitative statistical mechanics of relaxation kinetics.

### 6.3 Strengths and Limitations

The primary strength of this framework lies in its analytical rigour, its ability to bridge clinical oncology with non-equilibrium statistical mechanics, and its demonstrated structural stability under severe parametric uncertainty (Table 4). By translating qualitative biological affinities into dimensionless transport rates, we establish a parameter-free modelling framework that offers a quantitative operator-theoretic perspective on the classical *seed-and-soil* [20] hypothesis through spectral relaxation. Identifying the lung and liver as topological sinks [29] exhibiting negligible dispersion under Monte Carlo [33] perturbations provides a possible physical interpretation for their ubiquitous late-stage involvement, independent of local microenvironmental variations [60]. Despite these strengths, the current formulation operates under specific physical and empirical boundaries. Physically, treating the host architecture as a quenched, static background neglects continuous-time co-evolutionary feedback loops, such as pre-metastatic niche formation or treatment-induced vascular remodelling [16]. Empirically, a major limitation stems from the structural opacity and stringent regulatory constraints governing official clinical databases. While population-based registries (such as NCI-SEER [56, 67]) and post-mortem series provide robust cohort-level cross-sectional data, strict ethical and privacy frameworks (e.g., GDPR and HIPAA) [32] severely restrict access to longitudinal, patient-level micro-data. Consequently, operator-theoretic approaches operating at the ensemble topological level currently represent the most analytically tractable strategy for systemic risk stratification under existing data-governance boundaries.

### 6.4 Future Directions

The physical assumptions and data boundaries of the present framework point toward three principal avenues for future research in non-equilibrium statistical mechanics:

- Non-Markovian memory and anomalous transport: The present model assumes a first-order Markovian process [21], where transition probabilities depend solely on the current state [44]. Incorporating history-dependent dynamics, organ residence times, or non-Markovian memory retention [7] will require extending the operator framework to Generalised Langevin Equations [53] or fractional calculus diffusion equations [38].
- Annealed and adaptive higher-order landscapes: Transitioning from a quenched, static host architecture to an adaptive network landscape remains a major theoretical challenge. Future models must incorporate time-dependent transition rates Λ*_ij_*(*t*) and non-linear feedback loops where propagating disease dynamically remodels host connectivity through angiogenesis, niche priming, or therapeutic perturbations [16].
- Privacy-preserving micro-data integration: To bridge ensemble spectral predictions with individual clinical trajectories without violating data-privacy frameworks, future work should explore federated learning and synthetic patient cohort generation [66]. Coupling these computational strategies with continuous-time Markov chains could enable multi-scale validation at single-patient resolution as trajectory-resolved histories become accessible.

### 6.5 Concluding Remarks

Our findings indicate that, within the assumptions of the model, systemic metastatic dissemination tends to converge toward a stationary distribution in which initial biological preferences become progressively modulated by higher-order host connectivity patterns. The robustness of this behaviour under extensive parametric perturbations suggests that the predicted occupancy profiles are primarily determined by global network organisation rather than by isolated transition parameters [39]. From an operator-theoretic perspective, the present framework provides a quantitative interpretation of two relevant features of metastatic dissemination. First, it describes the dual functional role of lymph nodes (17.839% asymptotic probability mass), describing their transition from early dissemination intermediates (*n* = 1) toward highly connected compartments that redistribute metastatic flux across distal visceral organs in the non-equilibrium steady state. Second, the characteristic relaxation time (*τ* ≈ 1.49 discrete steps, Eq. 14) indicates that, according to the model dynamics, information associated with the initial primary site becomes progressively attenuated after a limited number of dissemination transitions (*n* ≥ 2, consistent with multi-organ metastatic progression patterns such as M1b/M1c disease). These results suggest that network-based descriptions could complement current tumour-centric approaches by highlighting recurrent host-level attractors, including lung, liver, and major nodal basins, as potential targets for risk stratification beyond primary tumour histology.

Taken together, the findings summarised in Table 5 support a host-network perspective of metastatic progression in which systemic vulnerability emerges from the interplay between tumour-derived dissemination tendencies and the structural organisation of the host environment. Although direct therapeutic modulation of network connectivity remains speculative, identifying persistent topological sinks and transport bottlenecks may provide a conceptual basis for future intervention strategies [36]. More broadly, the present operator-theoretic framework establishes a quantitative foundation for future models incorporating continuous-time clinical data, adaptive network dynamics, and physiological spectral representations [62]. Taken together, these findings support the view that higher-order host-network organisation constitutes a fundamental determinant of long-term metastatic transport, complementing tumour-centred descriptions with a quantitative systems-level perspective.

## Acknowledgements

This work is financed by the Multiannual Research Projects (PIP) N°11220200100439CO of the Consejo Nacional de Investigaciones Científicas y Técnicas (CONICET), Argentina. Funded by the European Union. Views and opinions expressed are however those of the author(s) only and do not necessarily reflect those of the European Union or European Research Executive Agency (REA) (granting authority). Neither the European Union nor the granting authority can be held responsible for them. The project leading to this application has received funding from the European Union’s Horizon Europe programme under the MSCA-SE grant agreement N°101131463 - SIMBAD: Statistical Inference from Multiscale Biological Data: Theory, Algorithms, Applications.

## Footnotes

1 Both *τ* and *τ*_log_ are bijective functions of the same sub-dominant eigenvalue *|λ*_2_*| ≈* 0.33. Consequently, the underlying operational dynamics, transition probabilities, and invariant attractor remain strictly invariant under either timescale convention (see Appendix B of the Supplementary Material for explicit proof).

