## Supplementary material for "Markovian Dynamics and Spectral Relaxation of Metastatic Networks"

#### Contents

**A. Structural Matrices and Row-Stochastic Transport Operators** **2**

**B. Spectral Decomposition and Asymptotic Approximations** **4**

**C. Computational Algorithms and Numerical Work-flows** **5**

### A. Structural Matrices and Row-Stochastic Transport Operators

This section presents the explicit mathematical representations of the primary matrices parameterising the metastatic transport model. These include the unweighted structural adjacency matrix $H$, which maps fundamental topological connections; the propensity operator $\Lambda$ and its corresponding weighted incidence matrix $H_w$, which incorporate clinical affinity weights; the transition matrix $W$, capturing raw directional fluxes; and the row-stochastic transition matrix $\tilde{W}$, which enforces global probability conservation and governs the discrete-time Markovian dynamics across the hypergraph backbone.

The unweighted structural adjacency matrix $H \in \{0,1\}^{m \times k}$, where $H_{ij}$ indicates the presence (1) or absence (0) of a topological connection from primary tumour $j$ to metastatic site $i$:

$H =$

| | Head & neck | Lung | Kidney | Pancreatic | Bladder | Testicular | Melanoma | Uterine | Rectal | Prostate | Thyroid | Oesophageal | Breast | Stomach | Colon | Ovarian | Liver | Cervical | STS | Ocular | BS |
|---|---|---|---|---|---|---|---|---|---|---|---|---|---|---|---|---|---|---|---|---|---|
| Lymph nodes | 1 | 1 | 0 | 0 | 1 | 1 | 1 | 1 | 0 | 1 | 1 | 1 | 1 | 1 | 0 | 1 | 1 | 1 | 0 | 0 | 0 |
| Bone | 1 | 1 | 1 | 1 | 1 | 1 | 1 | 1 | 1 | 1 | 1 | 1 | 1 | 1 | 1 | 1 | 1 | 1 | 1 | 1 | 0 |
| Brain | 1 | 1 | 1 | 1 | 1 | 1 | 1 | 0 | 1 | 1 | 1 | 1 | 1 | 0 | 1 | 1 | 1 | 1 | 1 | 1 | 1 |
| Pleura | 0 | 1 | 0 | 0 | 1 | 0 | 0 | 0 | 0 | 0 | 0 | 0 | 1 | 0 | 0 | 1 | 0 | 0 | 0 | 0 | 0 |
| Diaphragm | 0 | 1 | 0 | 0 | 0 | 0 | 0 | 0 | 0 | 0 | 0 | 0 | 0 | 0 | 0 | 1 | 0 | 0 | 0 | 0 | 0 |
| Liver | 1 | 1 | 0 | 1 | 1 | 1 | 1 | 1 | 1 | 1 | 1 | 1 | 1 | 1 | 1 | 1 | 0 | 1 | 1 | 1 | 1 |
| Kidney | 0 | 1 | 0 | 0 | 1 | 0 | 0 | 0 | 0 | 1 | 0 | 0 | 0 | 0 | 0 | 0 | 0 | 0 | 0 | 0 | 1 |
| Adrenal | 0 | 1 | 1 | 0 | 1 | 0 | 1 | 0 | 1 | 1 | 0 | 1 | 1 | 0 | 0 | 0 | 1 | 0 | 1 | 0 | 1 |
| Thyroid | 0 | 1 | 1 | 0 | 0 | 0 | 0 | 0 | 0 | 0 | 0 | 0 | 0 | 0 | 0 | 0 | 0 | 0 | 0 | 0 | 0 |
| Stomach | 0 | 0 | 0 | 1 | 0 | 0 | 0 | 0 | 0 | 0 | 0 | 0 | 0 | 0 | 1 | 0 | 0 | 0 | 0 | 0 | 0 |
| Colon | 0 | 0 | 0 | 1 | 1 | 0 | 0 | 0 | 0 | 0 | 0 | 0 | 0 | 0 | 0 | 1 | 0 | 0 | 0 | 0 | 0 |
| Peritoneum | 0 | 1 | 0 | 1 | 1 | 0 | 1 | 1 | 1 | 1 | 1 | 0 | 1 | 1 | 1 | 1 | 1 | 1 | 0 | 0 | 0 |
| Rectum | 0 | 0 | 0 | 0 | 1 | 0 | 0 | 0 | 0 | 0 | 0 | 0 | 0 | 0 | 0 | 0 | 0 | 0 | 0 | 0 | 0 |
| Prostate | 0 | 0 | 0 | 0 | 1 | 0 | 0 | 0 | 0 | 0 | 0 | 0 | 0 | 0 | 0 | 0 | 0 | 0 | 0 | 0 | 0 |
| Ureter | 0 | 0 | 0 | 0 | 1 | 0 | 0 | 0 | 0 | 0 | 0 | 0 | 0 | 0 | 0 | 0 | 0 | 0 | 0 | 0 | 0 |
| Lung | 1 | 0 | 1 | 1 | 1 | 1 | 1 | 1 | 1 | 1 | 1 | 1 | 1 | 1 | 1 | 1 | 1 | 1 | 1 | 1 | 1 |
| Ovary | 0 | 0 | 0 | 0 | 0 | 0 | 0 | 1 | 0 | 0 | 0 | 0 | 0 | 1 | 0 | 0 | 0 | 0 | 0 | 0 | 0 |
| Bladder | 0 | 0 | 0 | 0 | 0 | 0 | 0 | 0 | 0 | 0 | 0 | 0 | 0 | 0 | 1 | 0 | 0 | 0 | 0 | 0 | 0 |
| Skin | 1 | 1 | 1 | 0 | 0 | 0 | 0 | 0 | 0 | 0 | 0 | 0 | 1 | 0 | 1 | 1 | 0 | 0 | 0 | 1 | 0 |
| Pancreas | 0 | 1 | 1 | 0 | 0 | 0 | 1 | 0 | 0 | 1 | 0 | 0 | 1 | 0 | 0 | 0 | 0 | 0 | 0 | 0 | 0 |
| Spleen | 0 | 1 | 1 | 0 | 0 | 0 | 1 | 0 | 0 | 0 | 0 | 0 | 1 | 0 | 1 | 0 | 0 | 0 | 0 | 0 | 0 |
| Ocular | 0 | 0 | 0 | 0 | 0 | 0 | 0 | 0 | 1 | 0 | 0 | 0 | 1 | 0 | 1 | 0 | 0 | 0 | 0 | 0 | 0 |
| Breast | 0 | 1 | 1 | 0 | 0 | 0 | 0 | 0 | 0 | 0 | 0 | 0 | 0 | 0 | 0 | 0 | 0 | 0 | 0 | 0 | 0 |

*Note: Soft tissue sarcoma and bone sarcoma are abbreviated as STS and BS, respectively. These abbreviations are used in matrix $H$, as well as in $\Lambda$ and $H_w$.*

The propensity operator $\Lambda$, where $\Lambda_{ij} \in [0,1]$ represents the clinical affinity weight quantifying the preference of primary cancer $j$ to metastasise to organ $i$:

$\Lambda =$

| | Head & neck | Lung | Kidney | Pancreatic | Bladder | Testicular | Melanoma | Uterine | Rectal | Prostate | Thyroid | Oesophageal | Breast | Stomach | Colon | Ovarian | Liver | Cervical | STS | Ocular | BS |
|---|---|---|---|---|---|---|---|---|---|---|---|---|---|---|---|---|---|---|---|---|---|
| Lymph nodes | 1.0 | 1.0 | 0 | 0 | 0.1 | 1.0 | 1.0 | 0.1 | 0 | 1.0 | 0.1 | 1.0 | 1.0 | 0.1 | 0 | 0.1 | 1.0 | 1.0 | 0 | 0 | 0 |
| Bone | 0.1 | 1.0 | 0.1 | 0.1 | 1.0 | 0.01 | 1.0 | 0.01 | 0.1 | 1.0 | 1.0 | 0.1 | 1.0 | 0.1 | 0.01 | 0.01 | 1.0 | 0.1 | 0.1 | 0.1 | 0 |
| Brain | 0.01 | 1.0 | 0.01 | 0.01 | 0.01 | 0.01 | 1.0 | 0 | 0.01 | 0.01 | 0.01 | 0.01 | 0.1 | 0 | 0.01 | 0.01 | 0.01 | 0.01 | 0.01 | 0.01 | 0.01 |
| Pleura | 0 | 0.1 | 0 | 0 | 0.01 | 0 | 0 | 0 | 0 | 0 | 0 | 0 | 0.1 | 0 | 0 | 0.1 | 0 | 0 | 0 | 0 | 0 |
| Diaphragm | 0 | 0.01 | 0 | 0 | 0 | 0 | 0 | 0 | 0 | 0 | 0 | 0 | 0 | 0 | 0 | 1.0 | 0 | 0 | 0 | 0 | 0 |
| Liver | 0.1 | 1.0 | 0.1 | 1.0 | 1.0 | 0.1 | 1.0 | 0.1 | 1.0 | 0.1 | 0.01 | 1.0 | 1.0 | 1.0 | 1.0 | 1.0 | 0 | 0.1 | 0.1 | 1.0 | 0.1 |
| Kidney | 0 | 0.01 | 0 | 0 | 0.01 | 0 | 0 | 0 | 0 | 0.01 | 0 | 0 | 0 | 0 | 0 | 0 | 0 | 0 | 0 | 0 | 0.01 |
| Adrenal | 0 | 1.0 | 0.01 | 0 | 0.1 | 0 | 0.01 | 0 | 0.01 | 0.01 | 0 | 0.01 | 0.1 | 0 | 0 | 0 | 0.1 | 0 | 0.01 | 0 | 0.01 |
| Thyroid | 0 | 0.01 | 0.01 | 0 | 0 | 0 | 0 | 0 | 0 | 0 | 0 | 0 | 0 | 0 | 0 | 0 | 0 | 0 | 0 | 0 | 0 |
| Stomach | 0 | 0 | 0 | 0.01 | 0 | 0 | 0 | 0 | 0 | 0 | 0 | 0 | 0 | 0 | 0.01 | 0 | 0 | 0 | 0 | 0 | 0 |
| Colon | 0 | 0 | 0 | 0.01 | 0.01 | 0 | 0 | 0 | 0 | 0 | 0 | 0 | 0 | 0 | 0 | 0.01 | 0 | 0 | 0 | 0 | 0 |
| Peritoneum | 0 | 0.01 | 0 | 1.0 | 0.1 | 0 | 0.1 | 0.01 | 0.1 | 0.01 | 0.01 | 0 | 0.01 | 1.0 | 0.1 | 1.0 | 1.0 | 0.1 | 0 | 0 | 0 |
| Rectum | 0 | 0 | 0 | 0 | 0.01 | 0 | 0 | 0 | 0 | 0 | 0 | 0 | 0 | 0 | 0 | 0 | 0 | 0 | 0 | 0 | 0 |
| Prostate | 0 | 0 | 0 | 0 | 0.01 | 0 | 0 | 0 | 0 | 0 | 0 | 0 | 0 | 0 | 0 | 0 | 0 | 0 | 0 | 0 | 0 |
| Ureter | 0 | 0 | 0 | 0 | 0.01 | 0 | 0 | 0 | 0 | 0 | 0 | 0 | 0 | 0 | 0 | 0 | 0 | 0 | 0 | 0 | 0 |
| Lung | 0.1 | 0 | 1.0 | 0.1 | 1.0 | 1.0 | 1.0 | 1.0 | 1.0 | 0.1 | 1.0 | 1.0 | 1.0 | 0.1 | 0.1 | 0.01 | 1.0 | 1.0 | 1.0 | 0.1 | 1.0 |
| Ovary | 0 | 0 | 0 | 0 | 0 | 0 | 0 | 0.1 | 0 | 0 | 0 | 0 | 0 | 0.01 | 0 | 0 | 0 | 0 | 0 | 0 | 0 |
| Bladder | 0 | 0 | 0 | 0 | 0 | 0 | 0 | 0 | 0 | 0 | 0 | 0 | 0 | 0 | 0.01 | 0 | 0 | 0 | 0 | 0 | 0 |
| Skin | 0.01 | 0.01 | 0.01 | 0 | 0 | 0 | 0 | 0 | 0 | 0 | 0 | 0 | 0.1 | 0 | 0.01 | 0.01 | 0 | 0 | 0 | 0.01 | 0 |
| Pancreas | 0 | 0.01 | 0.01 | 0 | 0 | 0 | 0.01 | 0 | 0 | 0.01 | 0 | 0 | 0.01 | 0 | 0 | 0 | 0 | 0 | 0 | 0 | 0 |
| Spleen | 0 | 0.01 | 0.01 | 0 | 0 | 0 | 0.01 | 0 | 0 | 0 | 0 | 0 | 0.01 | 0 | 0.01 | 0 | 0 | 0 | 0 | 0 | 0 |
| Ocular | 0 | 0 | 0 | 0 | 0 | 0 | 0 | 0 | 0.01 | 0 | 0 | 0 | 0.01 | 0 | 0.01 | 0 | 0 | 0 | 0 | 0 | 0 |
| Breast | 0 | 0.01 | 0.01 | 0 | 0 | 0 | 0 | 0 | 0 | 0 | 0 | 0 | 0 | 0 | 0 | 0 | 0 | 0 | 0 | 0 | 0 |

The weighted incidence matrix $H_w = H \odot \Lambda$ combines structural connectivity with clinical propensity via the Hadamard (entrywise) product:

$H_w =$

| | Head & neck | Lung | Kidney | Pancreatic | Bladder | Testicular | Melanoma | Uterine | Rectal | Prostate | Thyroid | Oesophageal | Breast | Stomach | Colon | Ovarian | Liver | Cervical | STS | Ocular | BS |
|---|---|---|---|---|---|---|---|---|---|---|---|---|---|---|---|---|---|---|---|---|---|
| Lymph nodes | 1.0 | 1.0 | 0 | 0 | 0.1 | 1.0 | 1.0 | 0.1 | 0 | 1.0 | 0.1 | 1.0 | 1.0 | 0.1 | 0 | 0.1 | 1.0 | 1.0 | 0 | 0 | 0 |
| Bone | 0.1 | 1.0 | 0.1 | 0.1 | 1.0 | 0.01 | 1.0 | 0.01 | 0.1 | 1.0 | 1.0 | 0.1 | 1.0 | 0.1 | 0.01 | 0.01 | 1.0 | 0.1 | 0.1 | 0.1 | 0 |
| Brain | 0.01 | 1.0 | 0.01 | 0.01 | 0.01 | 0.01 | 1.0 | 0 | 0.01 | 0.01 | 0.01 | 0.01 | 0.1 | 0 | 0.01 | 0.01 | 0.01 | 0.01 | 0.01 | 0.01 | 0.01 |
| Pleura | 0 | 0.1 | 0 | 0 | 0.01 | 0 | 0 | 0 | 0 | 0 | 0 | 0 | 0.1 | 0 | 0 | 0.1 | 0 | 0 | 0 | 0 | 0 |
| Diaphragm | 0 | 0.01 | 0 | 0 | 0 | 0 | 0 | 0 | 0 | 0 | 0 | 0 | 0 | 0 | 0 | 1.0 | 0 | 0 | 0 | 0 | 0 |
| Liver | 0.1 | 1.0 | 0.1 | 1.0 | 1.0 | 0.1 | 1.0 | 0.1 | 1.0 | 0.1 | 0.01 | 1.0 | 1.0 | 1.0 | 1.0 | 1.0 | 0 | 0.1 | 0.1 | 1.0 | 0.1 |
| Kidney | 0 | 0.01 | 0 | 0 | 0.01 | 0 | 0 | 0 | 0 | 0.01 | 0 | 0 | 0 | 0 | 0 | 0 | 0 | 0 | 0 | 0 | 0.01 |
| Adrenal | 0 | 1.0 | 0.01 | 0 | 0.1 | 0 | 0.01 | 0 | 0.01 | 0.01 | 0 | 0.01 | 0.1 | 0 | 0 | 0 | 0.1 | 0 | 0.01 | 0 | 0.01 |
| Thyroid | 0 | 0.01 | 0.01 | 0 | 0 | 0 | 0 | 0 | 0 | 0 | 0 | 0 | 0 | 0 | 0 | 0 | 0 | 0 | 0 | 0 | 0 |
| Stomach | 0 | 0 | 0 | 0.01 | 0 | 0 | 0 | 0 | 0 | 0 | 0 | 0 | 0 | 0 | 0.01 | 0 | 0 | 0 | 0 | 0 | 0 |
| Colon | 0 | 0 | 0 | 0.01 | 0.01 | 0 | 0 | 0 | 0 | 0 | 0 | 0 | 0 | 0 | 0 | 0.01 | 0 | 0 | 0 | 0 | 0 |
| Peritoneum | 0 | 0.01 | 0 | 1.0 | 0.1 | 0 | 0.1 | 0.01 | 0.1 | 0.01 | 0.01 | 0 | 0.01 | 1.0 | 0.1 | 1.0 | 1.0 | 0.1 | 0 | 0 | 0 |
| Rectum | 0 | 0 | 0 | 0 | 0.01 | 0 | 0 | 0 | 0 | 0 | 0 | 0 | 0 | 0 | 0 | 0 | 0 | 0 | 0 | 0 | 0 |
| Prostate | 0 | 0 | 0 | 0 | 0.01 | 0 | 0 | 0 | 0 | 0 | 0 | 0 | 0 | 0 | 0 | 0 | 0 | 0 | 0 | 0 | 0 |
| Ureter | 0 | 0 | 0 | 0 | 0.01 | 0 | 0 | 0 | 0 | 0 | 0 | 0 | 0 | 0 | 0 | 0 | 0 | 0 | 0 | 0 | 0 |
| Lung | 0.1 | 0 | 1.0 | 0.1 | 1.0 | 1.0 | 1.0 | 1.0 | 1.0 | 0.1 | 1.0 | 1.0 | 1.0 | 0.1 | 0.1 | 0.01 | 1.0 | 1.0 | 1.0 | 0.1 | 1.0 |
| Ovary | 0 | 0 | 0 | 0 | 0 | 0 | 0 | 0.1 | 0 | 0 | 0 | 0 | 0 | 0.01 | 0 | 0 | 0 | 0 | 0 | 0 | 0 |
| Bladder | 0 | 0 | 0 | 0 | 0 | 0 | 0 | 0 | 0 | 0 | 0 | 0 | 0 | 0 | 0.01 | 0 | 0 | 0 | 0 | 0 | 0 |
| Skin | 0.01 | 0.01 | 0.01 | 0 | 0 | 0 | 0 | 0 | 0 | 0 | 0 | 0 | 0.1 | 0 | 0.01 | 0.01 | 0 | 0 | 0 | 0.01 | 0 |
| Pancreas | 0 | 0.01 | 0.01 | 0 | 0 | 0 | 0.01 | 0 | 0 | 0.01 | 0 | 0 | 0.01 | 0 | 0 | 0 | 0 | 0 | 0 | 0 | 0 |
| Spleen | 0 | 0.01 | 0.01 | 0 | 0 | 0 | 0.01 | 0 | 0 | 0 | 0 | 0 | 0.01 | 0 | 0.01 | 0 | 0 | 0 | 0 | 0 | 0 |
| Ocular | 0 | 0 | 0 | 0 | 0 | 0 | 0 | 0 | 0.01 | 0 | 0 | 0 | 0.01 | 0 | 0.01 | 0 | 0 | 0 | 0 | 0 | 0 |
| Breast | 0 | 0.01 | 0.01 | 0 | 0 | 0 | 0 | 0 | 0 | 0 | 0 | 0 | 0 | 0 | 0 | 0 | 0 | 0 | 0 | 0 | 0 |

The raw transition matrix $W = H_w H_w^T - \mathrm{diag}(H_w H_w^T)$, where $W_{ij}$ captures the directional flux strength from source organ $i$ to destination organ $j$:

$W =$

| | Lymph nodes | Bone | Brain | Pleura | Diaphragm | Liver | Kidney | Adrenal | Thyroid | Stomach | Colon | Peritoneum | Rectum | Prostate | Ureter | Lung | Ovary | Bladder | Skin | Pancreas | Spleen | Ocular | Breast |
|---|---|---|---|---|---|---|---|---|---|---|---|---|---|---|---|---|---|---|---|---|---|---|---|
| Lymph nodes | 0 | 7.43 | 2.19 | 0.31 | 1.01 | 7.51 | 0.03 | 1.33 | 0.01 | 0 | 0.02 | 3.35 | 0.01 | 0.01 | 0.01 | 9.31 | 0.11 | 0 | 0.13 | 0.04 | 0.03 | 0.01 | 0.01 |
| Bone | 9.50 | 0 | 2.25 | 0.31 | 1.01 | 11.71 | 0.03 | 1.36 | 0.02 | 0.02 | 0.03 | 4.55 | 0.01 | 0.01 | 0.01 | 12.61 | 0.11 | 0.01 | 0.16 | 0.05 | 0.05 | 0.03 | 0.02 |
| Brain | 9.30 | 7.83 | 0 | 0.31 | 1.01 | 10.71 | 0.04 | 1.37 | 0.02 | 0.02 | 0.03 | 3.54 | 0.01 | 0.01 | 0.01 | 12.51 | 0 | 0.01 | 0.16 | 0.05 | 0.05 | 0.03 | 0.02 |
| Pleura | 2.20 | 3.01 | 1.12 | 0 | 1.01 | 4 | 0.02 | 1.20 | 0.01 | 0 | 0.02 | 1.12 | 0.01 | 0.01 | 0.01 | 2.01 | 0 | 0 | 0.12 | 0.02 | 0.02 | 0.01 | 0.01 |
| Diaphragm | 1.10 | 1.01 | 1.01 | 0.20 | 0 | 2 | 0.01 | 1 | 0.01 | 0 | 0.01 | 1.01 | 0 | 0 | 0 | 0.01 | 0 | 0 | 0.02 | 0.01 | 0.01 | 0 | 0.01 |
| Liver | 8.50 | 6.94 | 2.25 | 0.31 | 1.01 | 0 | 0.04 | 1.27 | 0.02 | 0.02 | 0.03 | 3.55 | 0.01 | 0.01 | 0.01 | 12.61 | 0.11 | 0.01 | 0.16 | 0.05 | 0.05 | 0.03 | 0.02 |
| Kidney | 2.10 | 3 | 1.03 | 0.11 | 0.01 | 2.20 | 0 | 1.12 | 0.01 | 0 | 0.01 | 0.12 | 0.01 | 0.01 | 0.01 | 2.10 | 0 | 0 | 0.01 | 0.02 | 0.01 | 0 | 0.01 |
| Adrenal | 6.10 | 6.40 | 2.18 | 0.21 | 0.01 | 6.40 | 0.04 | 0 | 0.02 | 0 | 0.01 | 1.33 | 0.01 | 0.01 | 0.01 | 9.10 | 0 | 0 | 0.12 | 0.05 | 0.04 | 0.02 | 0.02 |
| Thyroid | 1 | 1.10 | 1.01 | 0.10 | 0.01 | 1.10 | 0.01 | 1.01 | 0 | 0 | 0 | 0.01 | 0 | 0 | 0 | 1 | 0 | 0 | 0.02 | 0.02 | 0.02 | 0 | 0.02 |
| Stomach | 0 | 0.11 | 0.02 | 0 | 0 | 2 | 0 | 0 | 0 | 0 | 0.01 | 1.10 | 0 | 0 | 0 | 0.20 | 0 | 0.01 | 0.01 | 0 | 0.01 | 0.01 | 0 |
| Colon | 0.20 | 1.11 | 0.03 | 0.11 | 1 | 3 | 0.01 | 0.10 | 0 | 0.01 | 0 | 2.10 | 0.01 | 0.01 | 0.01 | 1.11 | 0 | 0 | 0.01 | 0 | 0 | 0 | 0 |
| Peritoneum | 6.50 | 7.43 | 2.19 | 0.31 | 1.01 | 9.31 | 0.03 | 1.33 | 0.01 | 0.02 | 0.03 | 0 | 0.01 | 0.01 | 0.01 | 8.41 | 0.11 | 0.01 | 0.13 | 0.04 | 0.04 | 0.03 | 0.01 |
| Rectum | 0.10 | 1 | 0.01 | 0.01 | 0 | 1 | 0.01 | 0.10 | 0 | 0 | 0.01 | 0.10 | 0 | 0.01 | 0.01 | 1 | 0 | 0 | 0 | 0 | 0 | 0 | 0 |
| Prostate | 0.10 | 1 | 0.01 | 0.01 | 0 | 1 | 0.01 | 0.10 | 0 | 0 | 0.01 | 0.10 | 0.01 | 0 | 0.01 | 1 | 0 | 0 | 0 | 0 | 0 | 0 | 0 |
| Ureter | 0.10 | 1 | 0.01 | 0.01 | 0 | 1 | 0.01 | 0.10 | 0 | 0 | 0.01 | 0.10 | 0.01 | 0.01 | 0 | 1 | 0 | 0 | 0 | 0 | 0 | 0 | 0 |
| Lung | 8.50 | 6.94 | 1.26 | 0.21 | 1 | 10.81 | 0.03 | 0.37 | 0.01 | 0.02 | 0.03 | 4.54 | 0.01 | 0.01 | 0.01 | 0 | 0.11 | 0.01 | 0.15 | 0.04 | 0.04 | 0.03 | 0.01 |
| Ovary | 0.20 | 0.11 | 0 | 0 | 0 | 1.10 | 0 | 0 | 0 | 0 | 0 | 1.01 | 0 | 0 | 0 | 1.10 | 0 | 0 | 0 | 0 | 0 | 0 | 0 |
| Bladder | 0 | 0.01 | 0.01 | 0 | 0 | 1 | 0 | 0 | 0 | 0.01 | 0 | 0.10 | 0 | 0 | 0 | 0.10 | 0 | 0 | 0.01 | 0 | 0.01 | 0.01 | 0 |
| Skin | 3.10 | 2.32 | 1.15 | 0.30 | 1.01 | 5.20 | 0.01 | 1.11 | 0.02 | 0.01 | 0.01 | 1.12 | 0 | 0 | 0 | 2.31 | 0 | 0.01 | 0 | 0.03 | 0.04 | 0.02 | 0.02 |
| Pancreas | 4 | 4.10 | 2.12 | 0.20 | 0.01 | 3.20 | 0.02 | 1.13 | 0.02 | 0 | 0 | 0.13 | 0 | 0 | 0 | 3.10 | 0 | 0 | 0.12 | 0 | 0.04 | 0.01 | 0.02 |
| Spleen | 3 | 3.11 | 2.12 | 0.20 | 0.01 | 4.10 | 0.01 | 1.12 | 0.02 | 0.01 | 0 | 0.22 | 0 | 0 | 0 | 3.10 | 0 | 0.01 | 0.13 | 0.04 | 0 | 0.02 | 0.02 |
| Ocular | 1 | 1.11 | 0.12 | 0.10 | 0 | 3 | 0 | 0.11 | 0 | 0.01 | 0 | 0.21 | 0 | 0 | 0 | 2.10 | 0 | 0.01 | 0.11 | 0.01 | 0.02 | 0 | 0 |
| Breast | 1 | 1.10 | 1.01 | 0.10 | 0.01 | 1.10 | 0.01 | 1.01 | 0.02 | 0 | 0 | 0.01 | 0 | 0 | 0 | 1 | 0 | 0 | 0.02 | 0.02 | 0.02 | 0 | 0 |

The row-stochastic transition matrix $\tilde{W}$, obtained by normalising the rows of $W$ such that $\tilde{W}_{ij} = \frac{W_{ij}}{\sum_k W_{ik}}$ (strictly satisfying $\sum_j \tilde{W}_{ij} = 1$), governs the discrete-time Markovian dynamics across the organ network:

$\tilde{W} =$

| | Lymph nodes | Bone | Brain | Pleura | Diaphragm | Liver | Kidney | Adrenal | Thyroid | Stomach | Colon | Peritoneum | Rectum | Prostate | Ureter | Lung | Ovary | Bladder | Skin | Pancreas | Spleen | Ocular | Breast |
|---|---|---|---|---|---|---|---|---|---|---|---|---|---|---|---|---|---|---|---|---|---|---|---|
| Lymph nodes | 0 | 0.2261 | 0.0666 | 0.0094 | 0.0307 | 0.2285 | 0.0009 | 0.0405 | 0.0003 | 0 | 0.0006 | 0.1019 | 0.0003 | 0.0003 | 0.0003 | 0.2833 | 0.0033 | 0 | 0.0040 | 0.0012 | 0.0009 | 0.0003 | 0.0003 |
| Bone | 0.2166 | 0 | 0.0513 | 0.0071 | 0.0230 | 0.2670 | 0.0007 | 0.0310 | 0.0005 | 0.0005 | 0.0007 | 0.1037 | 0.0002 | 0.0002 | 0.0002 | 0.2875 | 0.0025 | 0.0002 | 0.0036 | 0.0011 | 0.0011 | 0.0007 | 0.0005 |
| Brain | 0.1977 | 0.1665 | 0 | 0.0066 | 0.0215 | 0.2277 | 0.0009 | 0.0291 | 0.0004 | 0.0004 | 0.0006 | 0.0753 | 0.0002 | 0.0002 | 0.0002 | 0.2659 | 0 | 0.0002 | 0.0034 | 0.0011 | 0.0011 | 0.0006 | 0.0004 |
| Pleura | 0.1381 | 0.1890 | 0.0703 | 0 | 0.0634 | 0.2511 | 0.0013 | 0.0753 | 0.0006 | 0 | 0.0013 | 0.0703 | 0.0006 | 0.0006 | 0.0006 | 0.1262 | 0 | 0 | 0.0075 | 0.0013 | 0.0013 | 0.0006 | 0.0006 |
| Diaphragm | 0.1482 | 0.1361 | 0.1361 | 0.0270 | 0 | 0.2695 | 0.0013 | 0.1348 | 0.0013 | 0 | 0.0013 | 0.1361 | 0 | 0 | 0 | 0.0013 | 0 | 0 | 0.0027 | 0.0013 | 0.0013 | 0 | 0.0013 |
| Liver | 0.2297 | 0.1875 | 0.0608 | 0.0084 | 0.0273 | 0 | 0.0011 | 0.0343 | 0.0005 | 0.0005 | 0.0008 | 0.0959 | 0.0003 | 0.0003 | 0.0003 | 0.3407 | 0.0030 | 0.0003 | 0.0043 | 0.0014 | 0.0014 | 0.0008 | 0.0005 |
| Kidney | 0.1766 | 0.2523 | 0.0866 | 0.0093 | 0.0008 | 0.1850 | 0 | 0.0942 | 0.0008 | 0 | 0.0008 | 0.0101 | 0.0008 | 0.0008 | 0.0008 | 0.1766 | 0 | 0 | 0.0008 | 0.0017 | 0.0008 | 0 | 0.0008 |
| Adrenal | 0.1901 | 0.1995 | 0.0680 | 0.0065 | 0.0003 | 0.1995 | 0.0012 | 0 | 0.0006 | 0 | 0.0003 | 0.0415 | 0.0003 | 0.0003 | 0.0003 | 0.2837 | 0 | 0 | 0.0037 | 0.0016 | 0.0012 | 0.0006 | 0.0006 |
| Thyroid | 0.1555 | 0.1711 | 0.1571 | 0.0156 | 0.0016 | 0.1711 | 0.0016 | 0.1571 | 0 | 0 | 0 | 0.0016 | 0 | 0 | 0 | 0.1555 | 0 | 0 | 0.0031 | 0.0031 | 0.0031 | 0 | 0.0031 |
| Stomach | 0 | 0.0316 | 0.0057 | 0 | 0 | 0.5747 | 0 | 0 | 0 | 0 | 0.0029 | 0.3161 | 0 | 0 | 0 | 0.0575 | 0 | 0.0029 | 0.0029 | 0 | 0.0029 | 0.0029 | 0 |
| Colon | 0.0227 | 0.1259 | 0.0034 | 0.0125 | 0.1134 | 0.3401 | 0.0011 | 0.0113 | 0 | 0.0011 | 0 | 0.2381 | 0.0011 | 0.0011 | 0.0011 | 0.1259 | 0 | 0 | 0.0011 | 0 | 0 | 0 | 0 |
| Peritoneum | 0.1758 | 0.2009 | 0.0592 | 0.0084 | 0.0273 | 0.2518 | 0.0008 | 0.0360 | 0.0003 | 0.0005 | 0.0008 | 0 | 0.0003 | 0.0003 | 0.0003 | 0.2274 | 0.0030 | 0.0003 | 0.0035 | 0.0011 | 0.0011 | 0.0008 | 0.0003 |
| Rectum | 0.0298 | 0.2976 | 0.0030 | 0.0030 | 0 | 0.2976 | 0.0030 | 0.0298 | 0 | 0 | 0.0030 | 0.0298 | 0 | 0.0030 | 0.0030 | 0.2976 | 0 | 0 | 0 | 0 | 0 | 0 | 0 |
| Prostate | 0.0298 | 0.2976 | 0.0030 | 0.0030 | 0 | 0.2976 | 0.0030 | 0.0298 | 0 | 0 | 0.0030 | 0.0298 | 0.0030 | 0 | 0.0030 | 0.2976 | 0 | 0 | 0 | 0 | 0 | 0 | 0 |
| Ureter | 0.0298 | 0.2976 | 0.0030 | 0.0030 | 0 | 0.2976 | 0.0030 | 0.0298 | 0 | 0 | 0.0030 | 0.0298 | 0.0030 | 0.0030 | 0 | 0.2976 | 0 | 0 | 0 | 0 | 0 | 0 | 0 |
| Lung | 0.2490 | 0.2033 | 0.0369 | 0.0062 | 0.0293 | 0.3166 | 0.0009 | 0.0108 | 0.0003 | 0.0006 | 0.0009 | 0.1330 | 0.0003 | 0.0003 | 0.0003 | 0 | 0.0032 | 0.0003 | 0.0044 | 0.0012 | 0.0012 | 0.0009 | 0.0003 |
| Ovary | 0.0568 | 0.0313 | 0 | 0 | 0 | 0.3125 | 0 | 0 | 0 | 0 | 0 | 0.2869 | 0 | 0 | 0 | 0.3125 | 0 | 0 | 0 | 0 | 0 | 0 | 0 |
| Bladder | 0 | 0.0079 | 0.0079 | 0 | 0 | 0.7937 | 0 | 0 | 0 | 0.0079 | 0 | 0.0794 | 0 | 0 | 0 | 0.0794 | 0 | 0 | 0.0079 | 0 | 0.0079 | 0.0079 | 0 |
| Skin | 0.1743 | 0.1304 | 0.0646 | 0.0169 | 0.0568 | 0.2923 | 0.0006 | 0.0624 | 0.0011 | 0.0006 | 0.0006 | 0.0630 | 0 | 0 | 0 | 0.1298 | 0 | 0.0006 | 0 | 0.0017 | 0.0022 | 0.0011 | 0.0011 |
| Pancreas | 0.2195 | 0.2250 | 0.1164 | 0.0110 | 0.0005 | 0.1756 | 0.0011 | 0.0620 | 0.0011 | 0 | 0 | 0.0071 | 0 | 0 | 0 | 0.1701 | 0 | 0 | 0.0066 | 0 | 0.0022 | 0.0005 | 0.0011 |
| Spleen | 0.1740 | 0.1804 | 0.1230 | 0.0116 | 0.0006 | 0.2378 | 0.0006 | 0.0650 | 0.0012 | 0.0006 | 0 | 0.0128 | 0 | 0 | 0 | 0.1798 | 0 | 0.0006 | 0.0075 | 0.0023 | 0 | 0.0012 | 0.0012 |
| Ocular | 0.1264 | 0.1403 | 0.0152 | 0.0126 | 0 | 0.3793 | 0 | 0.0139 | 0 | 0.0126 | 0 | 0.0265 | 0 | 0 | 0 | 0.2655 | 0 | 0.0013 | 0.0139 | 0.0013 | 0.0025 | 0 | 0 |
| Breast | 0.1555 | 0.1711 | 0.1571 | 0.0156 | 0.0016 | 0.1711 | 0.0016 | 0.1571 | 0.0031 | 0 | 0 | 0.0016 | 0 | 0 | 0 | 0.1555 | 0 | 0 | 0.0031 | 0.0031 | 0.0031 | 0 | 0 |

### B. Spectral Decomposition and Asymptotic Approximations

#### Determination of the Spectral Gap

To formally establish why the long-term relaxation of the metastatic probability vector $\bar{p}(n)$ is governed exclusively by the subdominant eigenvalue $\lambda_2$, we examine the algebraic structure of the real-valued but non-symmetric transition operator $\tilde{W}$.

Let $\sigma(\tilde{W}) = \{\lambda_1, \lambda_2, \ldots, \lambda_M\}$ denote the spectrum of the transition operator $\tilde{W}$, with its eigenvalues sorted in descending order of their absolute magnitudes such that $1 = \lambda_1 > |\lambda_2| \geq |\lambda_3| \geq \cdots \geq |\lambda_M|$. For our empirical baseline configuration, the subdominant eigenvalue is strictly real ($\lambda_2 \approx 0.33$). Consequently, the state vector $\bar{p}(n)$ at any discrete time step $n$ can be explicitly decomposed via projection onto the biorthogonal basis of left ($\bar{u}_k$) and right ($\bar{v}_k$) eigenvectors:

$$\bar{p}(n) = \bar{\pi} + \sum_{k=2}^{M} c_k \lambda_k^n \bar{u}_k$$

where $c_k = \langle \bar{p}(0), \bar{v}_k \rangle$ represents the initial expansion coefficients dictated by the primary tumour's location, and $\bar{\pi} = \bar{u}_1$ is the stationary distribution vector.

To isolate the dominant physical mechanism driving the dissolution of the transient phase, we factor out the leading subdominant eigenvalue $\lambda_2^n$ from the summation series:

$$\bar{p}(n) = \bar{\pi} + \lambda_2^n \left[ c_2 \bar{u}_2 + \sum_{k=3}^{M} c_k \left( \frac{\lambda_k}{\lambda_2} \right)^n \bar{u}_k \right]$$

By virtue of our strict magnitude ordering, the algebraic ratio satisfies:

$$\left| \frac{\lambda_k}{\lambda_2} \right| < 1 \quad \forall \quad k \geq 3$$

In the discrete-time asymptotic limit where $n \gg 1$, these scaling fractions contract exponentially. Taking the limit as $n$ advances reveals:

$$\lim_{n \to \infty} \left| \frac{\lambda_k}{\lambda_2} \right|^n = 0$$

Consequently, the contributions of all higher-order transient modes ($k \geq 3$) vanish at an accelerated rate compared to the first subdominant mode. This algebraic contraction leaves the system's state tracking an asymptotic trajectory driven solely by the slowest decaying mode:

$$\bar{p}(n) \approx \bar{\pi} + c_2 \lambda_2^n \bar{u}_2$$

This rigorous simplification justifies the definition of the spectral gap $\gamma = 1 - |\lambda_2|$ as the unique structural bottleneck governing the system's relaxation rate.

##### Formal Bounds and Equivalence of Relaxation Timescales

To formally reconcile continuous-time logarithmic mixing conventions with discrete spectral gaps, consider the geometric decay of the dominant transient mode $|\lambda_2|^n$. In logarithmic Markov chain theory, this contraction is expressed exponentially as:

$$|\lambda_2|^n = \exp\left(-\frac{n}{\tau_{\log}}\right), \quad \text{where} \quad \tau_{\log} = \frac{1}{\ln(1/|\lambda_2|)}.$$

By invoking the standard logarithmic inequality $1 - x \leq \ln(1/x)$, which holds strictly for any $x \in (0, 1)$, and letting $x = |\lambda_2|$, we establish:

$$1 - |\lambda_2| \leq \ln\left(\frac{1}{|\lambda_2|}\right) \implies \frac{1}{\ln(1/|\lambda_2|)} \leq \frac{1}{1 - |\lambda_2|} \implies \tau_{\log} \leq \tau.$$

This inequality proves that the linear spectral timescale $\tau = (1 - |\lambda_2|)^{-1} \approx 1.49$ discrete steps acts as a strict upper bound (a conservative linear envelope) to the asymptotic mixing time $\tau_{\log} \approx 0.90$ discrete steps.

Furthermore, because both $\tau$ and $\tau_{\log}$ are strictly monotonic, bijective functions of the underlying subdominant eigenvalue $|\lambda_2| \approx 0.33$, the trajectory vector $\bar{p}(n)$, the nodal transition probabilities, and the asymptotic attractor $\bar{\pi}$ remain strictly invariant under either convention. Adopting $\tau$ throughout this work thus provides a conservative, model-independent upper bound for the dissipation of initial primary seed information.

#### C. Computational Algorithms and Numerical Work-flows

To ensure full reproducibility of the mathematical model, this appendix details the numerical protocols used in our simulations. Algorithm 1 describes the complete computational pipeline for discrete-time Markovian dynamics on the metastatic hypergraph. It details the construction of the non-Hermitian transfer operator $\widetilde{W}$, the extraction of spectral relaxation metrics $(\gamma, \tau)$, and the iterative propagation of state probabilities together with Information Entropy and Total Variation Distance (TVD).

**Algorithm 1:** Markovian Metastatic Kinetics $(H, \Lambda, \mathbf{p}^{(0)}, N_{\max})$

**Input:** Incidence matrix $H \in \{0,1\}^{M\times N}$, affinity matrix $\Lambda \in \mathbb{R}_{\geq 0}^{M\times N}$, initial state $\mathbf{p}^{(0)}$, max steps $N_{\max}$

**Output:** Operator $\tilde{W}$, stationary state $\boldsymbol{\pi}$, relaxation parameters $(\gamma, \tau)$, kinetics $\{\mathbf{p}^{(n)}\}$, $\{\mathrm{TVD}(n)\}$, $\{S(n)\}$

`// Step 1: Construction of the Non-Hermitian Transfer Operator`

$H_w \leftarrow H \odot \Lambda$ ; `// Weighted hypergraph matrix via Hadamard product`

$W \leftarrow H_w H_w^\top$ ; `// Projection onto target organ space` $(M \times M)$

$W \leftarrow W - \mathrm{diag}(\mathrm{diag}(W))$ ; `// Removal of self-loops`

$\tilde{W} \leftarrow \mathrm{diag}(W\mathbf{1})^{-1} W$ ; `// Row-stochastic normalisation` ($\sum_j \tilde{W}_{ij} = 1$)

`// Step 2: Spectral Decomposition and Equilibrium State`

$[\boldsymbol{\lambda}, V] \leftarrow \mathrm{eigs}(\tilde{W}^\top)$ ; `// Compute eigenvalues and left eigenvectors`

$[\boldsymbol{\lambda}, V] \leftarrow \mathrm{sortAbsDescending}(\boldsymbol{\lambda}, V)$ ; `// Sort eigenvalues and permute eigenvector columns`

$\boldsymbol{\pi} \leftarrow \mathbf{v}_1 / \sum_i (\mathbf{v}_1)_i$ ; `// Stationary state (principal eigenvector for` $\lambda_1 = 1$)

$\gamma \leftarrow 1 - |\lambda_2|$ ; `// Spectral gap`

$\tau \leftarrow 1/\gamma$ ; `// Characteristic relaxation time`

`// Step 3: Markov Propagation, Relaxation Kinetics, and Entropy`

**for** $n \leftarrow 1$ **to** $N_{max}$ **do**

  $\mathbf{p}^{(n)} \leftarrow \mathbf{p}^{(0)} \tilde{W}^n$ ; `// State probability vector at step` $n$

  $\mathrm{TVD}(n) \leftarrow \frac{1}{2} \sum_{i=1}^{M} |\mathbf{p}_i^{(n)} - \pi_i|$ ; `// Total Variation Distance to stationary state`

  $S(n) \leftarrow -\sum_{i:\mathbf{p}_i^{(n)}>0} \mathbf{p}_i^{(n)} \log_2 \mathbf{p}_i^{(n)}$ ; `// Shannon information entropy (bits)`

**end**

**return** $\tilde{W}, \boldsymbol{\pi}, \gamma, \tau, \{\mathbf{p}^{(n)}\}, \{\mathrm{TVD}(n)\}, \{S(n)\}$

To assess the structural stability of the fast-mixing regime and test the invariance of the steady-state organ hierarchy against parametric noise, Algorithm 2 outlines the Monte Carlo Global Sensitivity Analysis (GSA). This routine applies uniform multiplicative perturbations ($\pm\epsilon$) across the discrete affinity levels ($\Lambda_{ij} \in \{10^0, 10^{-1}, 10^{-2}\}$), reconstructing the operator across $N_{\text{iter}}$ realizations to yield ensemble-averaged metrics and empirical 95% confidence intervals.

**Algorithm 2:** Global Sensitivity Analysis MC ($H, \Lambda_{\text{base}}, \epsilon, N_{\text{iter}}$)

**Input:** Binary incidence matrix $H \in \{0,1\}^{M\times N}$, base affinity matrix $\Lambda_{\text{base}} \in \mathbb{R}_{\geq 0}^{M\times N}$, perturbation noise $\epsilon$, iterations $N_{\text{iter}}$

**Output:** Mean relaxation time $\bar{\tau}$, standard deviation $\sigma_\tau$, mean stationary state $\bar{\boldsymbol{\pi}}$, dispersion $\boldsymbol{\sigma}_{\boldsymbol{\pi}}$, 95% confidence intervals $\text{CI}_{95\%}$

```
// Step 1: Topological Identification of Affinity Patterns
mask_common ← (Λ_base = 10^0) ;                    // Mask for strong affinity weights
mask_occasional ← (Λ_base = 10^{-1}) ;             // Mask for moderate affinity weights
mask_rare ← (Λ_base = 10^{-2}) ;                   // Mask for weak affinity weights
// Step 2: Monte Carlo Parametric Sampling and Spectral Extraction
for i ← 1 to N_iter do
    V_c ~ 10^0 × U(1 − ε, 1 + ε) ;    // Sample element-wise common affinity perturbations
    V_o ~ 10^{-1} × U(1 − ε, 1 + ε) ;   // Sample element-wise occasional affinity perturbations
    V_r ~ 10^{-2} × U(1 − ε, 1 + ε) ;   // Sample element-wise rare affinity perturbations
    Λ^(i) ← 0_{M×N};
    Λ^(i)[mask_common] ← V_c;
    Λ^(i)[mask_occasional] ← V_o;
    Λ^(i)[mask_rare] ← V_r;
    H_w ← H ⊙ Λ^(i) ;                      // Reconstruct perturbed weighted matrix
    W ← H_w H_w^⊤ ;                        // Projection onto target organ space (M × M)
    W ← W − diag(diag(W)) ;                // Removal of self-loops
    W̃ ← diag(W1)^{-1} W ;                  // Row-stochastic normalisation
    [λ, V] ← eigs(W̃^⊤) ;                   // Spectral decomposition
    [λ, V] ← sortAbsDescending(λ, V) ;     // Sort eigenvalues and eigenvectors
    τ^(i) ← 1/(1 − |λ_2|) ;                // Relaxation time realisation i
    π^(i) ← v_1 / Σ_j (v_1)_j ;            // Stationary state realisation i
end
// Step 3: Statistical Aggregation and Interval Estimation
τ̄ ← mean({τ^(i)}), σ_τ ← std({τ^(i)}) ;              // Moments of relaxation time
π̄ ← mean({π^(i)}), σ_π ← std({π^(i)}) ;              // Moments of stationary distribution
CI_95% ← [percentile({π^(i)}, 2.5), percentile({π^(i)}, 97.5)] ;   // Empirical 95% bounds
return τ̄, σ_τ, π̄, σ_π, CI_95%
```
